# Discovery of microRNA-like RNAs during early fruiting body development in the model mushroom *Coprinopsis cinerea*

**DOI:** 10.1101/325217

**Authors:** Amy Yuet Ting Lau, Xuanjin Cheng, Chi Keung Cheng, Wenyan Nong, Man Kit Cheung, Raymond Hon-Fu Chan, Jerome Ho Lam Hui, Hoi Shan Kwan

**Author notes:** These authors contributed equally to this work (Amy Yuet Ting Lau, Xuanjin Cheng).

## Abstract

*Coprinopsis cinerea* is a model mushroom particularly suited to study fungal fruiting body development and the evolution of multicellularity in fungi. While microRNAs (miRNAs) are extensively studied in animals and plants for their essential roles in post-transcriptional regulation of gene expression, miRNAs in fungi are less well characterized and their potential roles in controlling mushroom development remain unknown. To identify miRNA-like RNAs (milRNAs) in *C. cinerea* and explore their expression patterns during the early developmental transition of mushroom development, small RNA libraries of vegetative mycelium and primordium were generated and putative milRNA candidates were identified following the standards of miRNA prediction in animals and plants. Two out of 22 novel predicted milRNAs, cci-milR-12c and cci-milR-13e-5p, were validated by northern blot and stem-loop reverse transcription real-time PCR. Cci-milR-12c was differentially expressed whereas the expression levels of cci-milR-13e-5p were similar in the two developmental stages. Target prediction of the validated milRNAs resulted in genes associated with fruiting body development, including pheromone, hydrophobin, cytochrome P450, and protein kinase. Besides, essential genes for miRNA biogenesis, including three coding for Dicer-like (DCL), two for Argonaute-like (AGO-like) and one for quelling deficient-2 (QDE-2) proteins, were identified in the *C. cinerea* genome. Phylogenetic analysis showed that the DCL and AGO-like proteins of *C. cinerea* were more closely related to those in other basidiomycetes and ascomycetes than to animals and plants. Taken together, our findings provided the first evidence of milRNAs in the model mushroom and their potential roles in regulating fruiting body development. Information on the evolutionary relationship of milRNA biogenesis proteins across kingdoms has also provided new insights into further functional and evolutionary studies of miRNAs.

## Introduction

MicroRNAs (miRNAs), about 22 nucleotides (nt) in length, are one of the major groups of functional small non-coding RNAs (ncRNAs) besides piwi-interacting RNAs (piRNAs) and short interfering RNAs (siRNAs) [1,2]. Previous studies have demonstrated the presence of miRNAs in nearly all eukaryotic lineages and their essential roles in various biological processes [3–9]. In plants, miRNAs play roles in tissue morphogenesis, stress response and stem development [4]. In animals, miRNAs regulate cell proliferation and differentiation, apoptosis, and different metabolic pathways during developmental transition [5–9]. MiRNAs mediate post-transcriptional gene silencing to regulate gene expression through base pairing their seed region (2-7 nt at the 5’-end) to the untranslated region (UTR) or opening reading frame of their target genes [1,10]. Plant miRNAs mediate mRNA cleavage through perfect complementarity to their targets. On the contrary, miRNAs in animals bind to their targets through imprecise complementarity [10]. The first miRNA-like RNA (milRNA) in filamentous fungi was described in *Neurospora crassa* in 2010, more than a decade later than in animals and plants [11]. Although milRNAs have been subsequently discovered in other fungi, such as *Sclerotinia sclerotiorum, Trichoderma reesei, Penicillium marneffei, Fusarium oxysporum*, and *Antrodia cinnamonmea*, the potential roles of milRNAs in the developmental processes of mushroom forming fungi are still largely unknown [12–16].

Fungi possess a fascinating morphological diversity, ranging from the simplest unicellular yeasts to macroscopic mushrooms [17]. *Coprinopsis cinerea,* commonly known as the ink cap, is one of the most morphologically complex fungi with a well-characterized genome [18]. *C. cinerea* is also a model mushroom that is commonly used to study the developmental processes in higher basidiomycete fungi. Under nutrient depletion and normal day-night rhythm, the undifferentiated vegetative mycelium of *C. cinerea* undergoes dynamic genetic and physiological changes to form a multihyphal structure, known as the fruiting body, through hyphal aggregation and mycelial differentiation [19–21]. Fruiting body development of *C. cinerea* is a rapid but complex process, consisting of six main stages: mycelium, initials, stage 1 and 2 primordium, young fruiting body, and mature fruiting body [22]. The whole process can be completed within two weeks when the fungus is cultured on artificial media with optimal conditions [22,23]. Understanding the molecular regulatory mechanisms during fruiting body initiation and development is one of the major goals of mycological studies. The most significant transcriptomic switch has been demonstrated during the transition from mycelium to primordium, which represents a developmental transition from a loose, undifferentiated structure to a compact and well-organized multicellular body plan [24, 25].

Regulatory roles of miRNAs have been demonstrated in various multicellular organisms. In fungi, the key components of RNA regulatory networks and stage-specific milRNAs have also been reported [9, 11, 16, 26–31]. Some miRNAs in animals and plants are expressed in a stage-specific or tissue-specific manner, suggesting their potential roles in maintaining tissue specificity and functions [9, 32–35]. Here, we hypothesized that miRNAs also regulate developmental transition in the mushroom forming fungus *C. cinerea.* Prediction and identification of milRNAs and their targets in *C. cinerea* are feasible based on the published genome sequence data and transcriptomic profiles of the early developmental transition in *C. cinerea* [18, 24, 25]. In this study, we used high-throughput small RNA (sRNA) sequencing to computationally identify 22 putative milRNA candidates in the mycelium and primordium stages of *C. cinerea.* Two milRNAs, namely cci-milR-12c and cci-milR-13e-5p, were validated using northern blot analysis and their expression levels were examined by stem-loop RT-qPCR in both developmental stages. One of the milRNA candidates, cci-milR-12c, was found differentially expressed. Genes encoding putative Dicer-like (DCL), Argonuate-like (AGO-like) and quelling deficient-2 (QDE-2) proteins were identified in the *C. cinerea* genome [18]. Our results have provided evidence for the presence of milRNAs in *C. cinerea*, revealed their potential targets, and demonstrated a differential expression of milRNAs during the early developmental stages of the model mushroom. Our study has facilitated the understanding of the diversified regulatory roles of milRNAs and the molecular mechanisms of fruiting body development in higher basidiomycete fungi.

## Materials and Methods

### *C. cinerea* strains and growth conditions

The *C. cinerea* strain used for the identification of milRNAs and core milRNA biogenesis proteins is a dikaryon, mated from the monokaryotic strains J6;5-4 and J6;5-5 [36]. Two monokaryons were generated from single spore isolates of a dikaryon that had been backcrossed with the reference strain Okayama 7#130 for five generations. Besides, the homokaryotic fruiting strain #326 *(A43mut B43mut pab1-1)* was used in the siRNA-mediated Dicer knockdown analysis [37]. The strains were cultured on YMD medium containing 0.4% yeast extract, 1% malt extract, and 0.4% glucose with Bacto agar. Mycelia were cultivated on agar plates at 37 °C for about 4-5 days until the mycelium grew over the whole agar surface and reached the edge of plates. Fruiting body formation was induced by incubating the mycelium culture at 25 °C under a light/dark regime of 12/12 h [18,38,39]. The incubator was kept at a relative humidity > 60% for the production of fruiting bodies.

### RNA isolation and sRNA sequencing

Samples were collected from two biological replicates for each developmental stage of *C. cinerea.* In brief, total RNA was extracted from mycelium (MYC) (4-5 days in the dark) and primodium (PRI) (~ 6-7 mm tall, 3 days in the light) using mirVana miRNA Isolation Kit (Ambion) and treated with TURBO DNA-free Kit (Ambion) according to the manufacturer’s instructions. Mycelia from four agar plates and 4-5 independent primordium structures were harvested and pooled to form one replicate. All samples were stored at −80 °C. Concentration and quality of RNA samples were checked using an Agilent 2100 Bioanalyzer. Using total RNA as the starting material, sRNA sequencing was performed by Macrogen (Korea) on a Hiseq 2500 platform (Illumina).

### Bioinformatics analysis of sRNAs and prediction of milRNA candidates

Raw sequence reads were filtered to remove low quality reads with a Phred score lower than 20, adaptor and primer sequences, and reads shorter than 18 nt (Macrogen, Seoul, Korea). High quality reads were then used to build a non-redundant dataset in which reads identical in length and identity were clustered (Macrogen, Seoul, Korea). Clean unique reads were searched on Rfam v.9.1 to identify other types of small ncRNAs, such as rRNA, tRNA and snRNA (Macrogen, Seoul, Korea) [40]. Since previous studies have identified miRNAs derived from other small ncRNAs, clean clustered reads with 18-30 nt, including those aligned to tRNAs and rRNAs, were subsequently mapped to the *C. cinerea* genome (NCBI assembly accession: AACS00000000) using Bowtie and only perfectly matched sRNA reads were selected for milRNA prediction (Macrogen, Seoul, Korea) [41].

For milRNA candidate prediction, short clean reads ranging from 18-30 nt were first aligned to miRBase v.21 to categorize known miRNAs [42]. Then, an in-house Perl program was developed to identify the novel miRNAs. Since *N. crassa* demonstrated a wider range of precursor length than that in plants, the remaining mapped sRNA sequences were first extended on the genome to 51-150 bp in length to form a precursor-like hairpin structure [11]. Secondary structures of the extended sequence were computed by RNAfold in Vienna RNA package 2.0 with GU wobble base pair allowed. Putative milRNA candidates were selected using the following criteria: (1) sRNA that formed a hairpin structure with minimum free energy (MFE) of folding ≤ - 20 kcal/mol; (2) a RANDFOLD p-value of the predicted secondary structures < 0.01; and (3) with at least four reads [43,44,45].

### Validation of milRNAs by northern blot analysis

Northern blot analysis of milRNA identification was performed according to the protocol of Kim et al. with double-labeled digoxigenin (DIG) oligonucleotide probes instead of locked nucleic acids (LNA) probes [46]. Briefly, total RNA (5-15 ug) from the two different developmental stages were resolved on a 15% denaturing polyacrylamide gel with 8M Urea in 1X TBE. The RNA gels were then transferred to Hybond-N+ (Amersham Biosciences) at 10-15 V (30-60 min) using a Trans-Blot SD semi-dry transfer cell (Bio-Rad). Cross-linking, hybridization and membrane detection were performed as previously described [46]. Cross-linking was performed using freshly prepared 1-ethyl-3-(3-dimethylaminopropyl) carbodiimide (EDC) reagent at 60 °C for 1 hr. Membranes were hybridized overnight in ULTRAhyb™ hybridization buffer (Ambion) with specific double DIG-labeled oligonucleotide probes synthesized by Integrated DNA Technologies at 37 °C. Sequences of the probes used against the putative milRNAs were as follows: ccin-milR-12c, 5’-AAAGGTAGTGGTATTTCAACGGCGCC-3’; ccin-milR-13e-5p, 5’-AGTCCCTACTAGGTCCCGAG-3’. Probe detection was performed using DIG luminescent detection kit according to the manufacturer’s instructions (Roche) and photoemissions were detected using the ChemiDoc-It Imaging System (Bio-rad).

### Identification and phylogenetic analysis of DCL and AGO protein genes

One DCL protein (XP_002911949.1), two AGO-like proteins (XP_001837237.2, XP_001837864.2) and a QDE-2 protein (XP_001838344.1) were found in the predicted protein sequences of *C. cinerea* from *GenBank* (AACS00000000) [18]. For the phylogenetic analyses of the two main effector proteins in miRNA biogenesis, Dicer and AGO, corresponding protein sequences of animals, plants and some ascomycete fungi were downloaded from UniProt (http://www.uniprot.org/). Based on the annotated protein sequences of *N. crassa* DCL-1, DCL-2 (XP_961898.1, XP_963538.3) and QDE-2 (XP_958586.1), two other DCL proteins of *C. cinerea* (XP_001837094.2, XP_001840952.1), some ascomycete and basidiomycete fungi were identified using BLASTP against the JGI database (https://genome.jgi.doe.gov/) [11]. An E-value of ≤ 10E-10 and an identity ≥ 25% were used as the cutoffs in the BLASTP searches. Functional domains of the corresponding proteins in *C. cinerea* were predicted using Pfam and SMART [47, 48]. Phylogenetic trees of the proteins were constructed by the maximum likelihood method with 1000 bootstrap replicates using MEGA 7 [49].

### Experimental quantification of milRNAs and biogenesis proteins by RT-qPCR

The sequence-specific TaqMan MicroRNA Assays and TaqMan small RNA Assays (Life Technologies) were used for RT-qPCR of cci-milR-12c and cci-milR-13e-5p, and the 5S rRNA (endogenous control), respectively. Reverse transcription was performed using TaqMan MicroRNA Reverse Transcription Kit (Applied Biosystems, Inc). Results from the 5S rRNA were used for normalization. cDNA was amplified in 20 uL reaction mixtures containing TaqMan Universal PCR Master Mix, no AmpErase UNG (Applied Biosystem) using standard qPCR conditions (95 °C for 10 min, followed by 40 cycles of 95 °C for 15 sec and 60 °C for 1 min) [50].

To examine the expression levels of the Dicer and AGO proteins in *C. cinerea*, total RNA was reverse transcribed to cDNA using Transcriptor First Strand cDNA Synthesis Kit (Roche Applied Science) with random hexamer primers. Real-time PCR analysis was performed using the SsoAdvanced Universal SYBR Green Supermix (Bio-rad), with 1 μl of 10 μM gene-specific forward and reverse primers (S1 Table). Thermal cycling was performed for 35-40 cycles with each cycle comprising polymerase activation at 95 °C for 30 sec, denaturation at 95 °C for 5-15 sec, and extension at 60 °C for 1 min. The relative expressions of DCL, AGO-like and QDE-2 proteins were normalized against 18S rRNA with forward primer (5′-GCCTGTTTGAGTGTCATTAAATTCTC-3′) and reverse primer (5′-CTGCAACCCCCACATCCA-3′). All the cycling reactions were performed in triplicate and the cycle threshold fluorescence data were recorded on an ABI 7500 Fast Real-Time PCR system (Applied Biosystems). The comparative Ct method (ΔΔCt) was exploited to calculate the relative expression levels of both validated milRNAs, DCLs, AGO-like and QDE-2 proteins. Statistical analysis was performed by Student’s t-tests. A P-value <0.05 was considered statistically significant.

### Dicer-like proteins knockdown mediated by siRNAs

At least two sequence-specific siRNA targeting separate regions for each DCL mRNA were transfected into the stipe of primordium twice using needle and syringe to enhance the efficiency and effectiveness of knockdown [51]. The 5’-3’ sequences of the sense and antisense strands of synthetic Stealth siRNA duplexes (Invitrogen) of three DCLs are shown in S2 Table. The gene silencing effects were optimized through direct transfection of 8 μM siRNA. Briefly, primordium were first treated with synthetic siRNAs and incubated at 25 °C for 24 h. Then, the transfected primordia were treated with the same concentration of siRNAs and incubated for another 24 h. After the double transfection, total RNA samples of the control groups, untreated primordium and unrelated transfection (primordium with RNase-free water injection), and DCL knockdown strains were harvested. The remaining gene expressions of knockdown strains compared with the control groups were measured by quantitative real-time PCR using primers listed in S1 Table. Primers were designed to detect sequences between the sites of siRNA directed cleavage or at the target site of siRNA.

### MilRNA target prediction and functional annotation

Since most miRNAs bind to the 3’-UTR of their target mRNAs to down-regulated their gene expressions, a database was constructed from the 1,000 bp downstream sequences of the stop codon of all genes in the *C. cinerea* genome for miRNA target prediction [18]. Since there are no general rules for the complementarity between fungal milRNAs and their targets and no prediction algorithms have been developed for fungi, three different types of software, PITA, miRanda and microTar, were used here to predict the potential targets of validated milRNAs in order to minimize false positive results [52–55]. Target genes were selected only when they were predicted by all of the three software.

Additional filtering steps were applied to select for putative targets with annotated biological functions using Gene Ontology (GO) terms, Eukaryotic Orthologous Groups (KOG) groups, KEGG orthologs (KO), and KEGG biological pathways [56,57,58]. Target genes that have been found to involve in fruiting body development were considered as putative targets [59]. GO terms of targets were assigned using BLAST2GO (version 2.4.2) with default parameters. KOG groups were assigned by RPS-BLAST (E-value cut-off of 1.00E-3). KO and KEGG biological pathways were assigned with the KEGG Automatic Annotation Server (KAAS) using all available fungal species as the representative gene set and the bidirectional best hits method (BBH). To further identify target mRNAs that likely interact with milRNA in vivo, annotated putative targets with similar expression patterns to the corresponding milRNAs between the two developmental stages were selected based on previously published microarray data of *C. cinerea* [25]. Functional targets with a fold change ≤ 0.5 and > 0.5 at MYC compared to PRI were considered as putative milRNA targets of cci-milR-12c and cci-milR-13e-5p, respectively.

## Results

### Identification of sRNAs in *C. cinerea* by high-throughput sequencing

The general features of sRNA species uncovered in MYC and PRI of *C. cinerea* are shown in Table 1. A total of 16,925,614 and 17,490,760 raw reads were obtained from MYC and PRI, respectively. A total of 1,354,235 and 1,379,040 unique sRNA reads (18-30 nt) were obtained from the MYC and PRI stages, respectively. A total of 152,835 and 135,648 rRNAs, and 15,890 and 12,280 tRNAs were included in the unique clean reads of the MYC and PRI samples, respectively. The major proportion of sRNA clean reads from both stages was 20-22 nt in length (Fig 1a) and had a strong preference for 5’ uracil (Fig 1b).

**Table 1.**
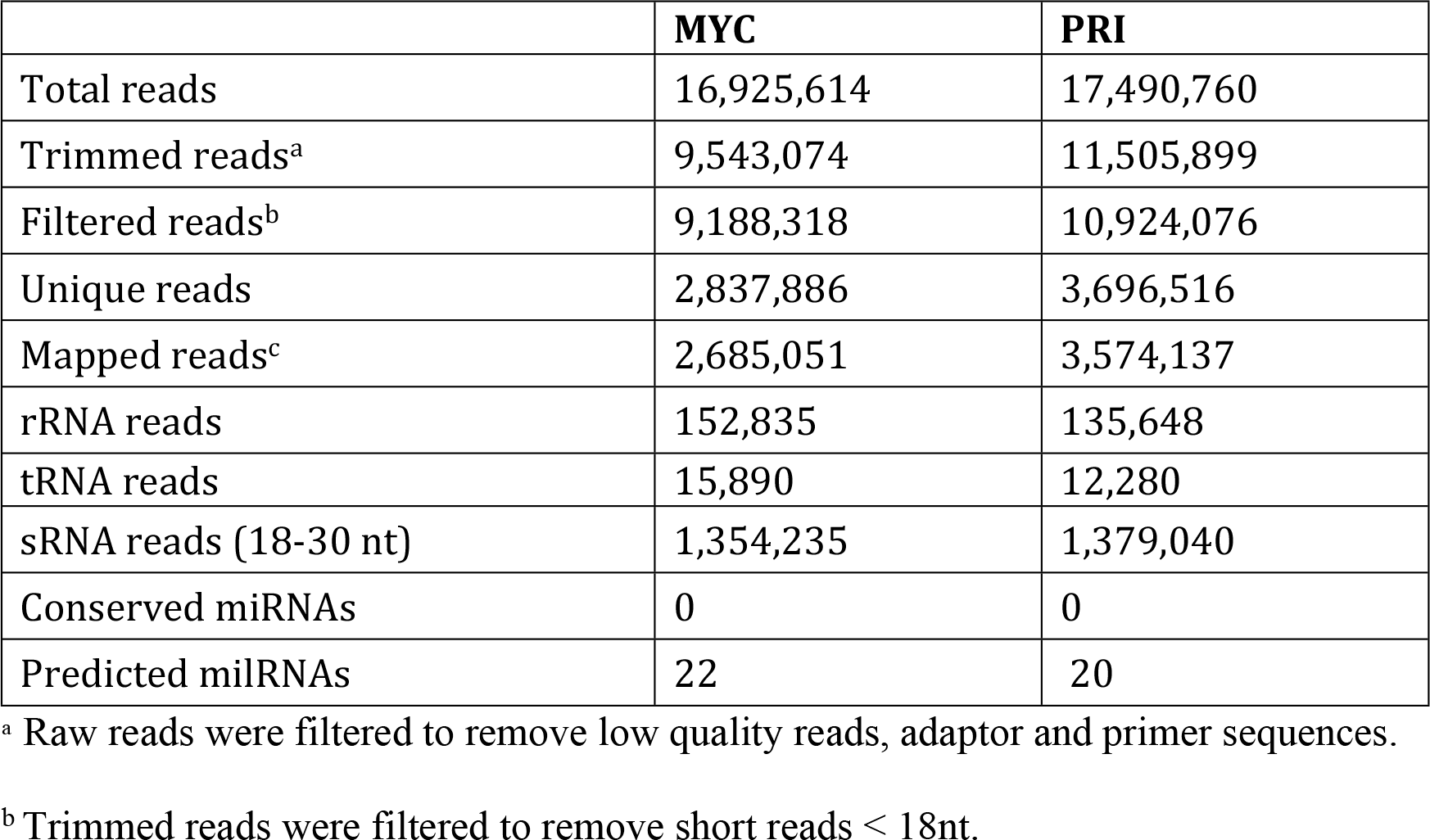
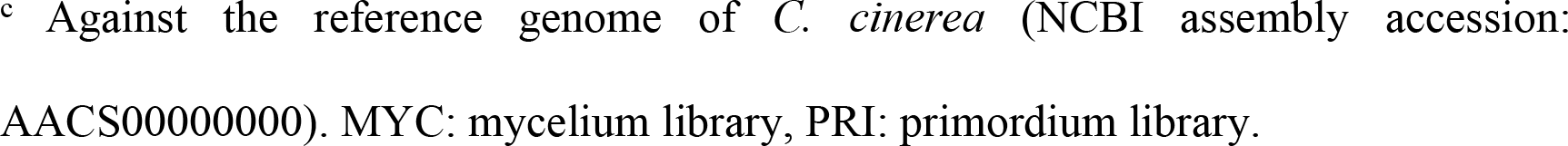
General features of sRNA sequencing of *C. cinerea* in two developmental stages.

|  | **MYC** | **PRI** |
| --- | --- | --- |
| Total reads | 16,925,614 | 17,490,760 |
| Trimmed reads<sup>a</sup> | 9,543,074 | 11,505,899 |
| Filtered reads<sup>b</sup> | 9,188,318 | 10,924,076 |
| Unique reads | 2,837,886 | 3,696,516 |
| Mapped reads<sup>c</sup> | 2,685,051 | 3,574,137 |
| rRNA reads | 152,835 | 135,648 |
| tRNA reads | 15,890 | 12,280 |
| sRNA reads (18-30 nt) | 1,354,235 | 1,379,040 |
| Conserved miRNAs | 0 | 0 |
| Predicted milRNAs | 22 | 20 |

<sup>a</sup> Raw reads were filtered to remove low quality reads, adaptor and primer sequences.
<sup>b</sup> Trimmed reads were filtered to remove short reads < 18nt.
° Against the reference genome of *C. cinerea* (NCBI assembly accession: AACSS000000000). MYC: mycelium library, PRI: primordium library.

**Fig 1.**
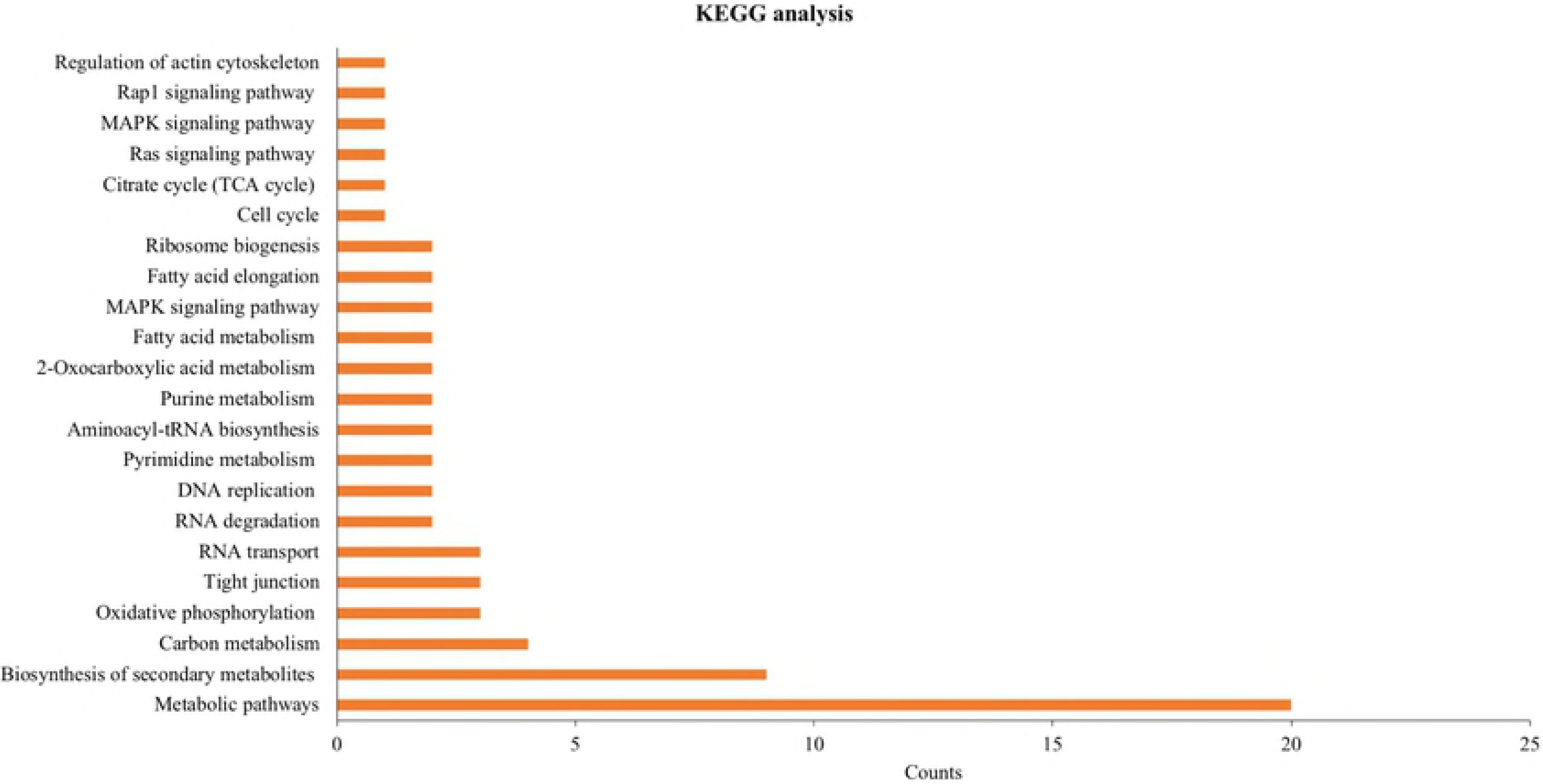
Features of sRNAs uncovered in *C. cinerea*. (a) Size distribution and (b) 5’ end nucleotide frequency of sRNAs in mycelial (MYC) and primordium (PRI) stages.

### Prediction and identification of potential milRNAs in *C. cinerea*

Twenty-two putative milRNA candidates were identified in *C. cinerea.* Most of them appeared in both MYC and PRI, except cci-milR-1 and cci-milR-2, which were only presented in MYC. The read counts and sequences of the milRNAs are listed in Table 2. The 20 and 26 nt classes were the most abundant groups in the milRNA candidates (Fig 2a). Guanine dominated the 5’ end nucleotide with weak superiority (Fig 2b). Similar to canonical miRNAs in animals and plants, most of the milRNAs in *C. cinerea* were derived from the intergenic region (68%), with five from rRNA (23%) and two from exon (9%) (Fig 2c). As for the gene locations, half of the putative milRNAs dwelled on the assembled chromosomes (Table 2). Six putative milRNAs predicted in *C. cinerea* were located within a short distance on the U413 contig, similar to milRNAs in animals and plants, which usually appear in clusters [60].

**Table 2.**
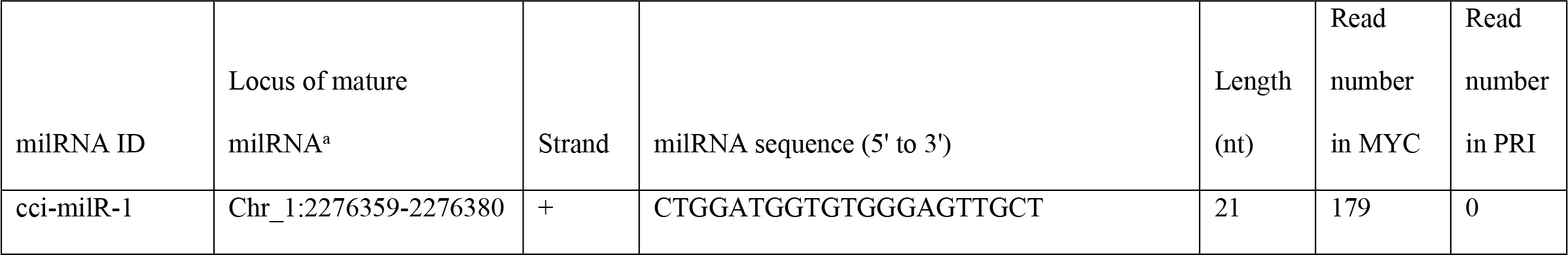
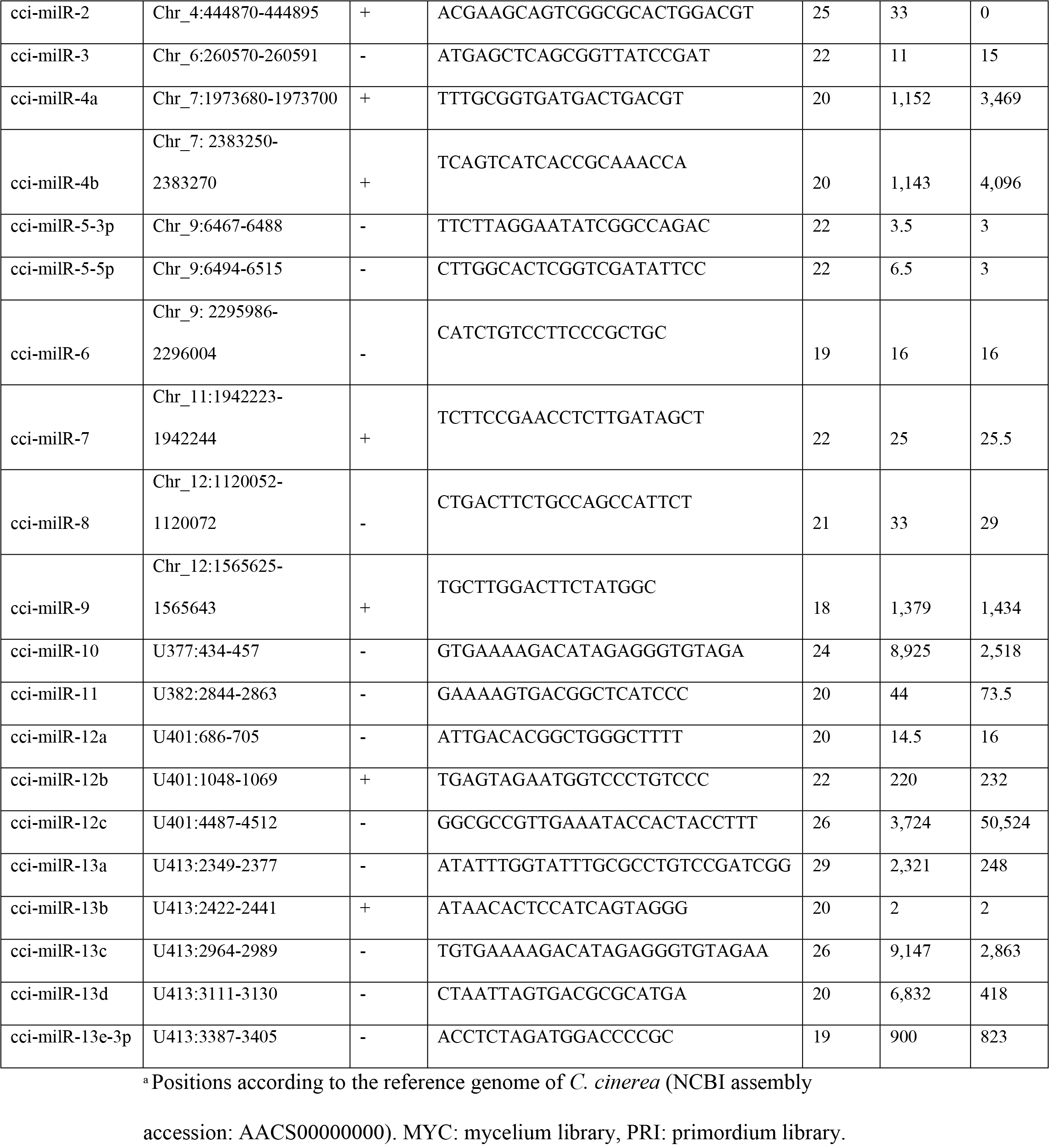
Twenty-two predicted milRNAs in *C. cinerea.*

**Fig 2.**
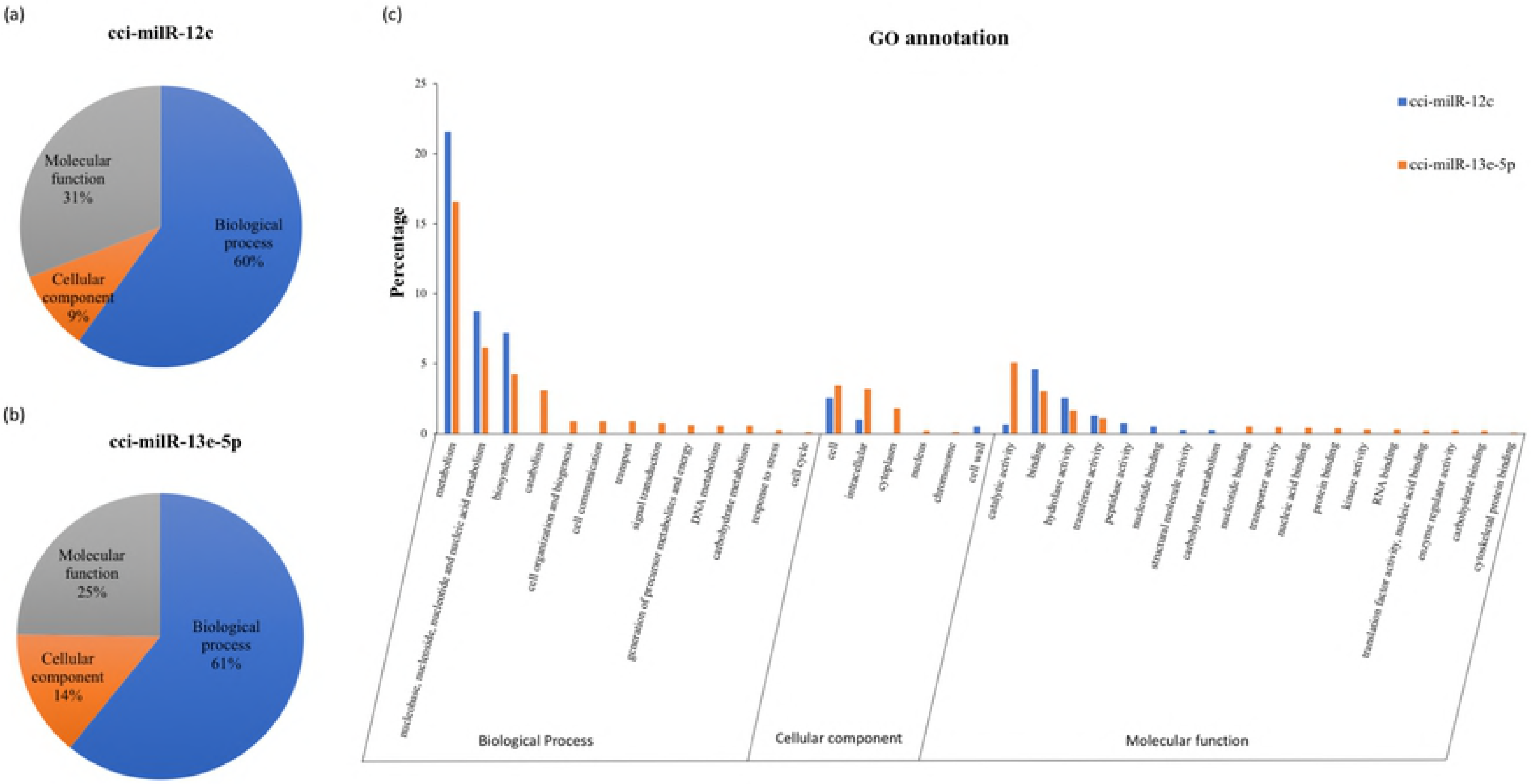
Characterization of putative milRNA candidates in *C. cinerea.* (a) Size distribution, (b) 5’ end nucleotide frequency and (c) annotation gene loci of the 22 putative milRNAs.

Although no conserved milRNA of animals and plants was found in *C. cinerea*, one homolog of cci-milR-12c was identified in another mushroom forming basidiomycete fungus, *Laccaria bicolor* (GSE9784), suggesting important regulatory roles of milRNAs in mushroom forming fungi. Sequence of the cci-milR-12c precursor (pre-milR-12c) was BLAST searched against the *L. bicolor* EST database [61], and only sequences with no mismatches on the seed region of mature cci-milR-12c and with less than three mismatches to the downstream sequence of the seed region were regarded as homologs [62]. The absence of milRNA homologs of animals and plants in *C. cinerea* indicates evolutionary divergence of miRNA genes among these three kingdoms, coinciding with most of the fungal milRNAs [12,13,16].

### Validation and characterization of milRNA expression patterns

Northern blot and RT-qPCR were used to validate the presence and to examine the expression levels of putative milRNAs in the two developmental stages. Two out of 22 milRNA candidates, cci-milR-12c and cci-milR-13e-5p, were verified using northern blot and their expression patterns during this developmental transition are shown in Fig 3. cci-milR-12c showed a higher expression in PRI (fold change >2), meaning that this milRNA candidate was differentially expressed in the early developmental stage. By contrast, the expression level of cci-milR-13e-5p in MYC was only slightly higher than that in PRI. The hairpin precursors of the two validated milRNAs are shown in Fig 4. Annotation of the gene loci of these two miRNAs indicated that the cci-milR-12c gene is located on an unassembled contig and cci-milR-13e-5p is derived from the intergenic region, based on the genome assembly data (AACS00000000) [17].

**Fig 3.**
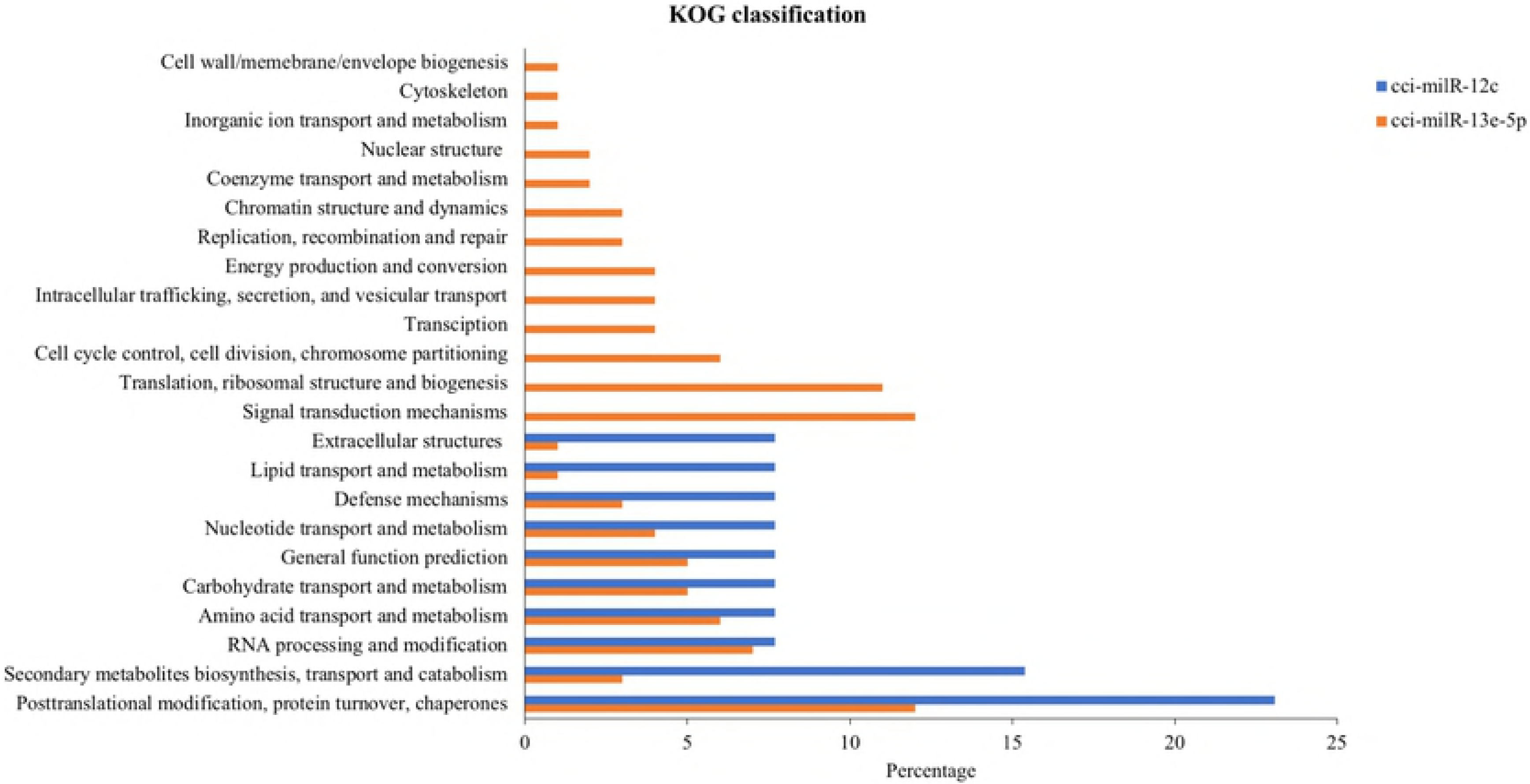
Validation of two milRNA candidates by northern blot and RT-qPCR. MYC: mycelium, PRI: primordium. Northern blot of sRNA samples shows the presence of (a) cci-milR-12c and (b) cci-milR-13e-5p in both developmental stages of *C. cinerea*. The top panel shows northern blots probed with the milRNA-specific DIG probes. The 15% denaturing gel stained with ethidium bromide (EtBr) in the bottom panel indicates equal loading of RNA samples. RT-qPCR results show the expression levels of (c) cci-milR-12c and (d) cci-milR-13e-5p in both developmental stages. Results were obtained from three independent experimental replicates and were significantly different between stages. ** p < 0.01, *** p< 0.001.

**Fig 4.**
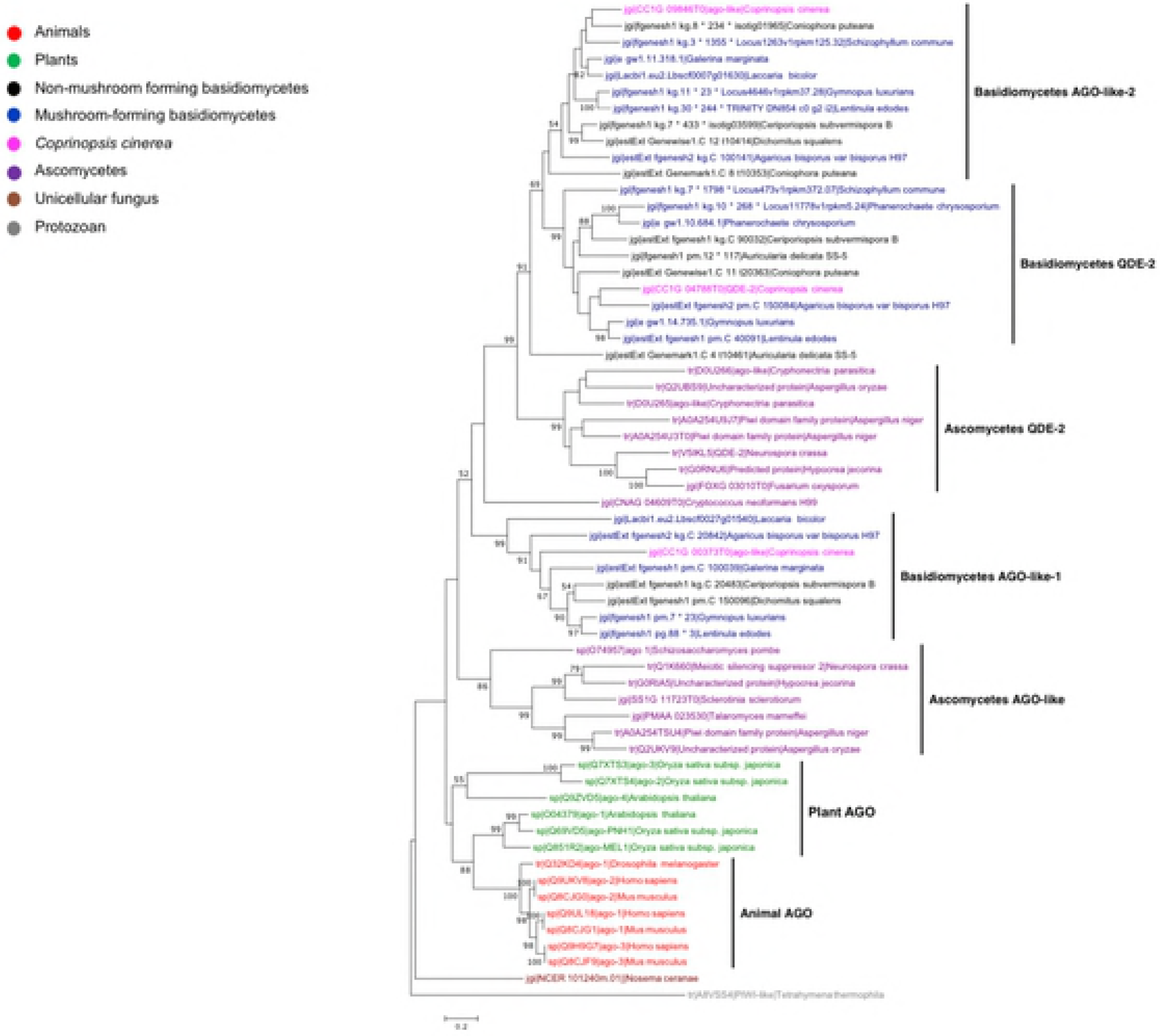
Predicted secondary structures of milRNA precursors. The predicted structures of pre-milR-12c and pre-milR-13e-5p with mature milRNA sequences labeled in red.

Although no annotated genes were identified in the locus of mature cci-milR-12c, there was a significant hit of pre-milR-12c to a eukaryotic rRNA sequence (accession: RF02543) when searching the precursor sequence on the Rfam database (E-value = 1.6e-26). Nucleotide sequence search also revealed that the pre-milR-12c sequence matched to the 28S rDNA locus of *C. cinerea* (E-value = 8e-62). Similar to rRNA-derived miRNAs found in human, the location of rRNA genes recovered here was distinct from that of the rRNA genes and cci-milR-12c might be generated during processing of the transcribed rRNA gene [63].

### Identification of DCL, AGO-like and QDE-2 proteins and characterization of their expression patterns in MYC and PRI

Dicer and AGO are effector proteins known to participate in miRNA biogenesis in animals and plants [64]. QDE-2 protein is an AGO that has been identified to involve in the pre-miRNA cleavage in *N. crassa* and its homologs have also been found in various fungal species [11, 14, 15]. Based on homolog search of the *N. crassa* DCL, AGO-like and QDE-2 proteins against the *C. cinerea genome* and annotated protein sequences from Broad Institute, three DCL, two AGO-like and a QDE-2 protein were found in *C. cinerea* (Fig 5) [11, 59]. These proteins were named DCL-1 (CC1G_00230), DCL-2 (CC1G_03181), DCL-3 (CC1G_13988), AGO-like-1 (CC1G_00373), AGO-like-2 (CC1G_09846), and QDE-2 (CC1G_04788). The AGO-like genes CC1G_00373 (3,979 bp in length) and CC1G_09846 (3,457 bp in length) encode mRNAs that result in 897 and 981 amino acid residues, respectively. By contrast, the QDE-2 gene (CC1G_04788) is 3,438 bp in length and its resultant mRNA generates a 965 amino acid residue. All AGO family proteins predicted in *C. cinerea* contained at least one of the two characteristic domains of AGO protein: PAZ and Piwi domain. The mRNAs of Dicer protein homologs (CC1G_00230, CC1G_03181, CC1G_13988) encode 1499, 2074 and 1457 amino acid residues, respectively. Interestingly, only one of the *C. cinerea* DCLs (CC1G_00230) contained the PAZ domain, which is present only in mushroom-specific DCLs but not other fungal DCLs. Although PAZ is a conserved domain in both Dicer and many AGO family proteins, it cannot be found in one of our annotated AGO-like proteins (CC1G_00373) in *C. cinerea* (Fig 5) [65,66]. In general, the domain organization of DCL, AGO-like and QDE-2 proteins of *C. cinerea* was similar to that of *N. crassa*, except that there is no PAZ domain in AGO-like-1 (S1 Fig).

**Fig 5.**
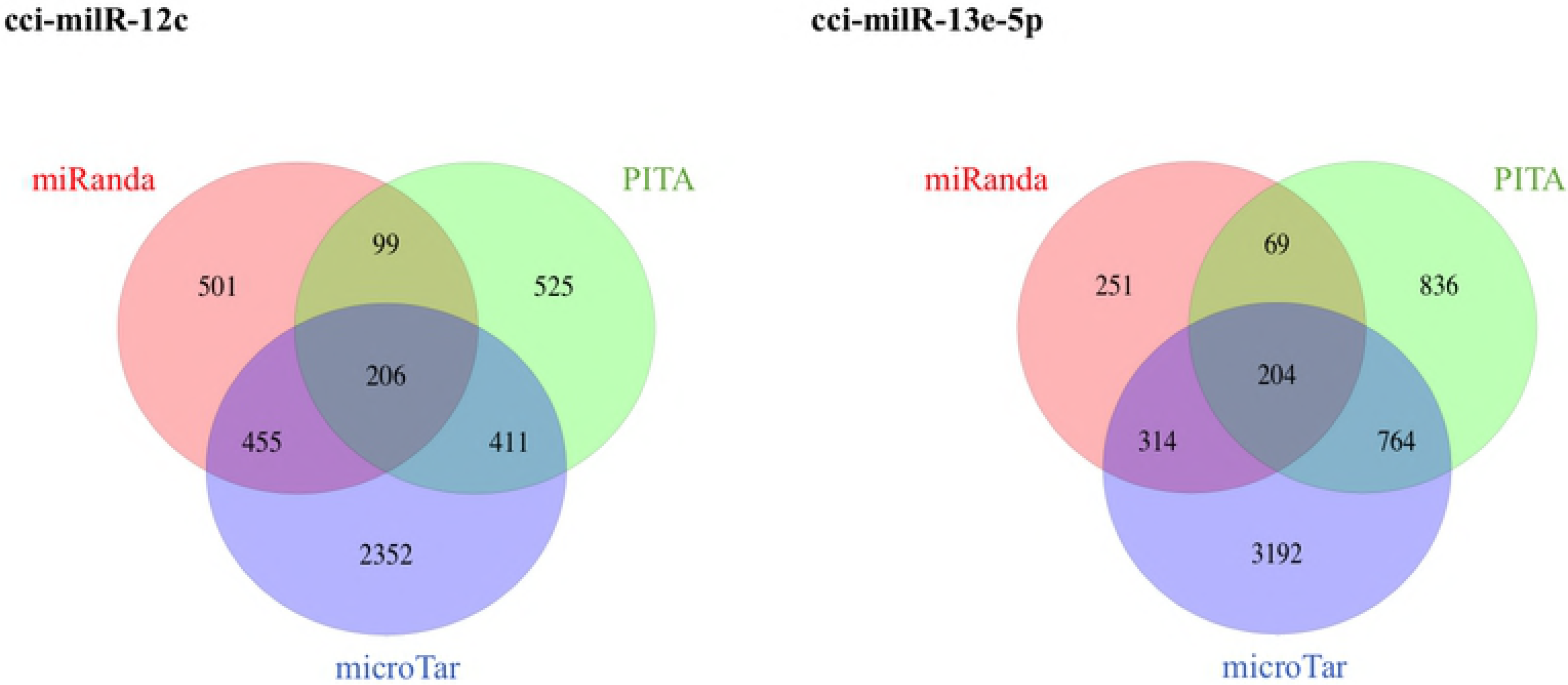
Schematic 2D domain architecture of Dicer and AGO proteins in *C. cinerea.* The grey bars represent the full protein sequences and the colored boxes represent identified functional domains.

RT-qPCR was used to examine the mRNA expression levels of the protein homologs and the results are shown in Fig 6. DCL-1 and DCL-2 showed higher expression levels in PRI, however, DCL-3 was down regulated in PRI (Fig 6a). For the AGO homologs, the expression levels of AGO-like-1 and QDE-2 were significantly lower in PRI (Fig 6b). Similar to the expression levels of cci-milR-12c, DCL-1 and DCL-2, AGO-like-2 was expressed significantly higher in PRI than MYC. Therefore, DCL-1 or DCL-2 and AGO-like-2 are more likely involved in the biogenesis of cci-milR-12c. On the contrary, AGO-like-1 or QDE-2 and DCL-3 are more likely related to the higher expression of cci-milR-13e-5p in MYC.

**Fig 6.**
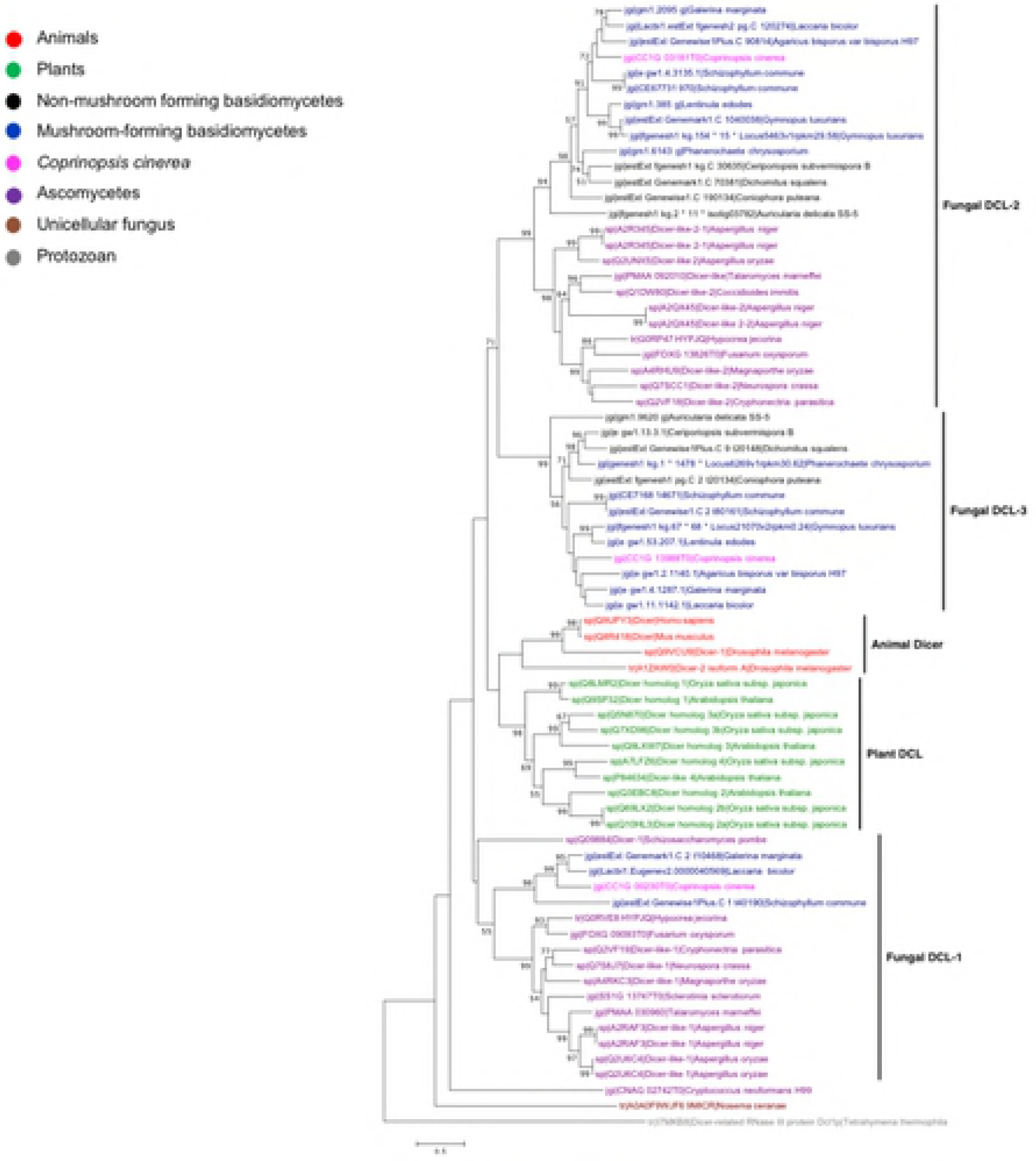
Relative mRNA expression levels of DCL and AGO proteins. The expression levels of (a) Dicer-like (DCLs) and (b) AGO-like and QDE-2 proteins in mycelium (MYC) and primordium (PRI) stages. Results were obtained from three independent experimental replicates and were significantly different between stages. *p < 0.05, ** p < 0.01, *** p< 0.001.

### Phylogenetic analysis of DCL and AGO homologs

Phylogenetic analysis of DCL and AGO proteins showed that both proteins duplicated early in the eukaryotic lineage and evolved independently in animals, plants and fungi (Figs 7 and 8). DCL and AGO homologs in *C. cinerea* were closely related to those in other basidiomycetes. Most of the mushroom forming fungi possess three DCLs, while other fungi contain only two DCLs. Besides, one DCL (CC1G_00230) of *C. cinerea* was grouped with DCLs of other mushroom forming basidiomycetes, namely *Galerina marginata, Laccaria bicolor* and *Schizophyllum commune* (Fig 7). These results suggest that Dicer proteins duplicated and diversified early in the eukaryotic lineage.

**Fig 7.**
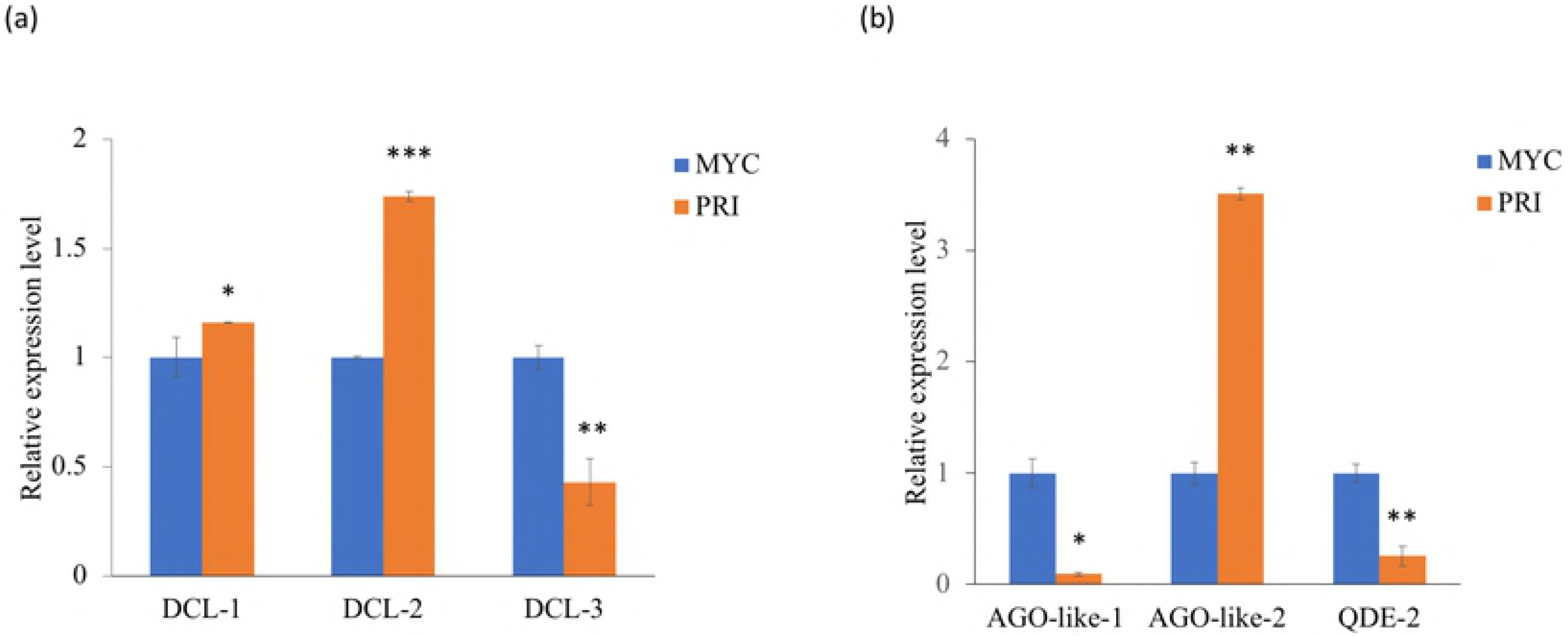
Phylogenetic tree of DCL proteins in animals, plants and fungi. The tree was constructed using the maximum likelihood method. Different groups of organisms were marked with different colors. The protozoan *Tetrahymena thermophile* was used as an outgroup. Bootstrap values were calculated from 1000 replicates and only values ≥50% were shown here. The scale bar represents 0.5 substitutions per nucleotide position.

**Fig 8.**
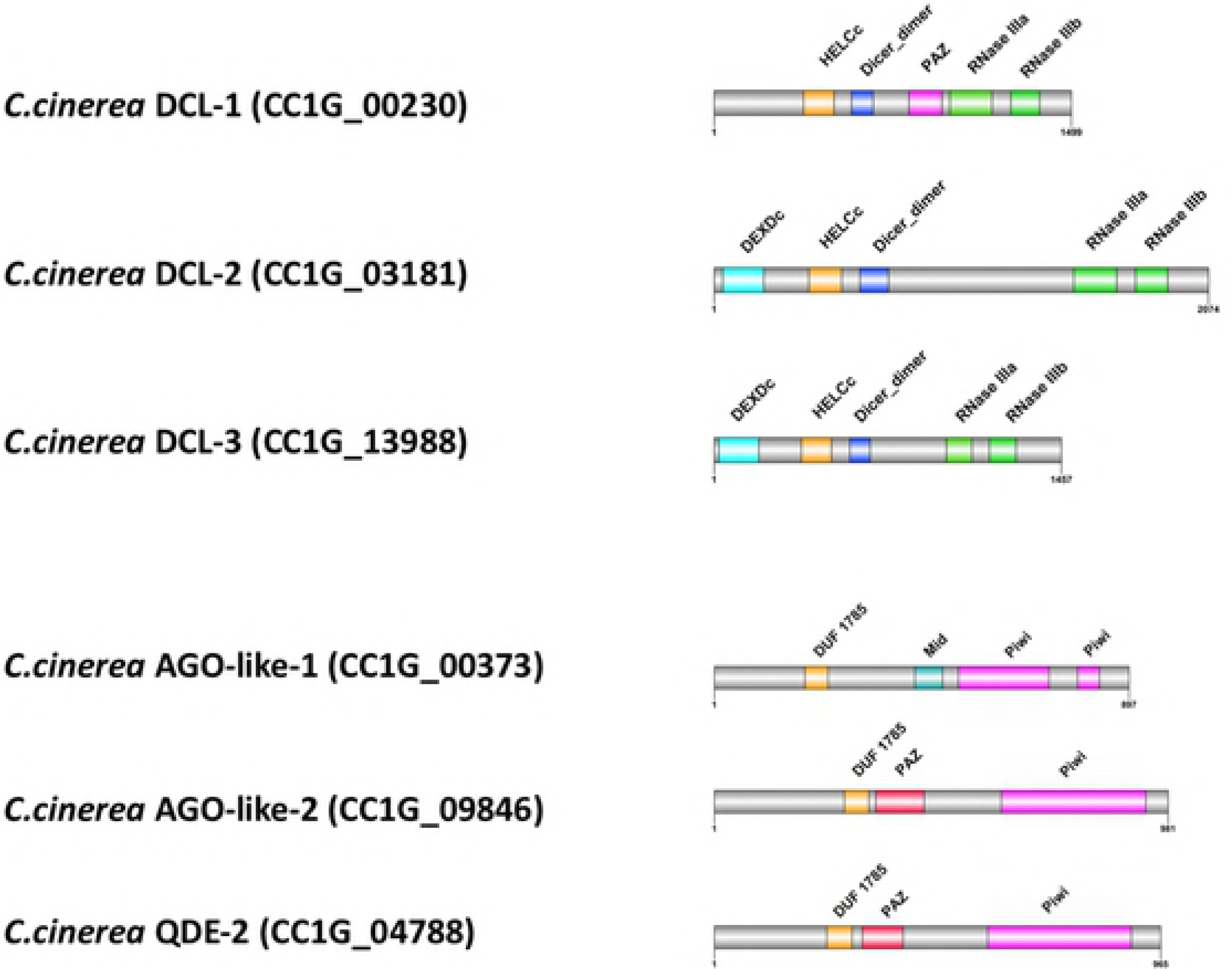
Phylogenetic tree of AGO proteins in animals, plants and fungi. The tree was constructed using the maximum likelihood method. Different groups of organisms were marked with different colors. The protozoan *Tetrahymena thermophile* was used as an outgroup. Bootstrap values were calculated from 1000 replicates and only values ≥50% were shown here. The scale bar represents 0.2 substitutions per nucleotide position.

### The expression levels of milRNAs in the DCL knockdown strains

Sequence-specific siRNA duplexes were used to knockdown individual DCL mRNA. The efficiency of knockdown of DCL mRNA following double transfection and the expression patterns of the two validated milRNAs in DCL knockdown strains are summarized in S2 Fig. Primordium transfected with siRNAs showed 60-80% knockdown of DCL mRNA transcript abundance comparing to the two control groups. A DIG-labelled probe specific for cci-milR-12c detected three bands in the control, with approximate sizes of 25/26, 40 and 50 nt on northern blot. The cci-milR-13e-5p-specific probe also revealed three bands on the blot, with sizes of about 20, 30, 40 nt. The ~20 nt bands were similar in size to the predicted cci-milR-12c and cci-milR-13e-5p, suggesting that they are the mature milRNAs. By constrast, the intermediate RNAs ~30-50 nt in size are likely the precursors of milRNAs (pre-milRNAs). However, the signals of the two mature milRNAs of DCL knockdown strains were similar to those of the controls. Given that the expression of DCL mRNAs was not completely abolished using knockdown, the roles of DCLs in milRNA biogenesis cannot be confirmed here.

### Prediction and functional annotation of milRNA targets

Computational prediction of milRNA targets was carried out based on three different algorithms: miRanda, PITA and microTar, to minimize false-positive results. Each prediction algorithm predicted a few hundreds to thousands of target genes for each milRNA. miRanda and PITA rely on evolutionary conservation to select functional targets whereas microTar discerns milRNA targets by calculating the duplex energies without taking into account the conservation of miRNA targets [52–56]. The number of overlapped targets is shown in a Venn diagram (Fig. 9). There were 206 and 204 common targets of cci-milR-12c and cci-milR-13e-5p, respectively. Of these, 143 and 140 were annotated with functional GO, KOG terms or fruiting body related genes (data not shown). Given that the expression patterns of milRNA are similar to their targets and two milRNAs showed higher expression in MYC and PRI respectively, the expression levels of putative targets during the transition from MYC to PRI were used for the last filtering step. As a result, 15 and 133 functional genes were selected as the putative targets of cci-milR-12c and cci-milR-13e-5p, respectively (S3 Table).

**Fig 9.**
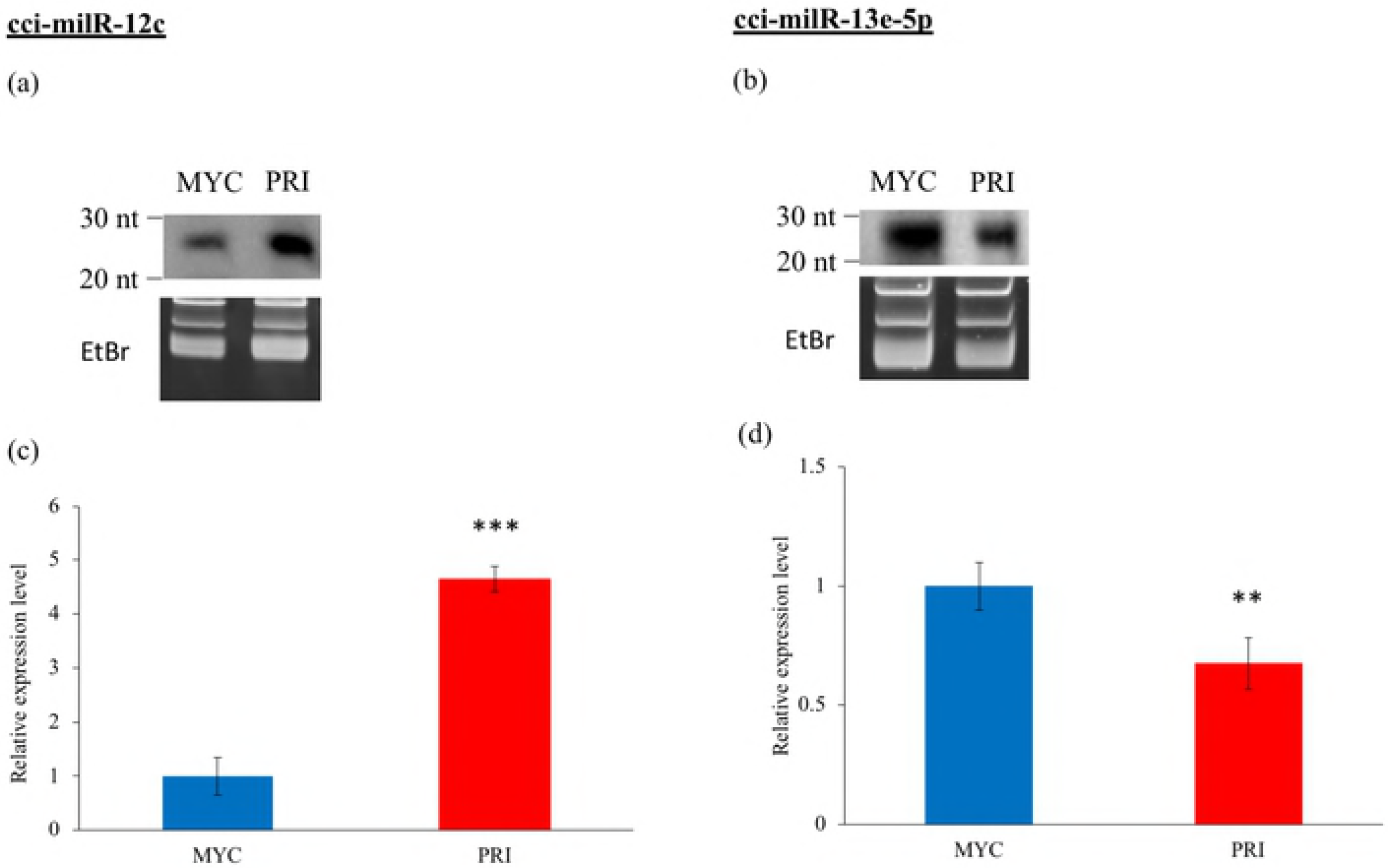
Venn diagram showing the distribution of the number of putative milRNA targets predicted by miRanda, PITA and microTar. To fully understand the functions of the putative targets of milRNA, the targets were annotated using GO terms, KOG terms and KEGG pathway. Results of GO term annotation revealed that the majority (> 60%) of the putative targets of milRNAs were categorized to the biological processes (Figs 10a and 10b). For both validated milRNAs, “metabolism”, “nucleobase, nucleoside, nucleotide and nucleic acid metabolism”, and “biosynthesis” were the most remarkably enriched GO terms under this category (Fig 10c). Additional functional annotation of putative milRNA targets was performed by searching the eukaryotic homologs in the KOG database. Putative targets of cci-milR-12c were assigned to only 10 groups and cci-milR-13e-5p were assigned to 23 groups (Fig 11). For the KOG classifications of cci-milR-12c putative targets, the category “posttranslational modification, protein turnover, chaperones” (23%) was the largest group, followed by the “secondary metabolites biosynthesis, transport and catabolism” (15%) category (Fig 11).

**Fig 10.**
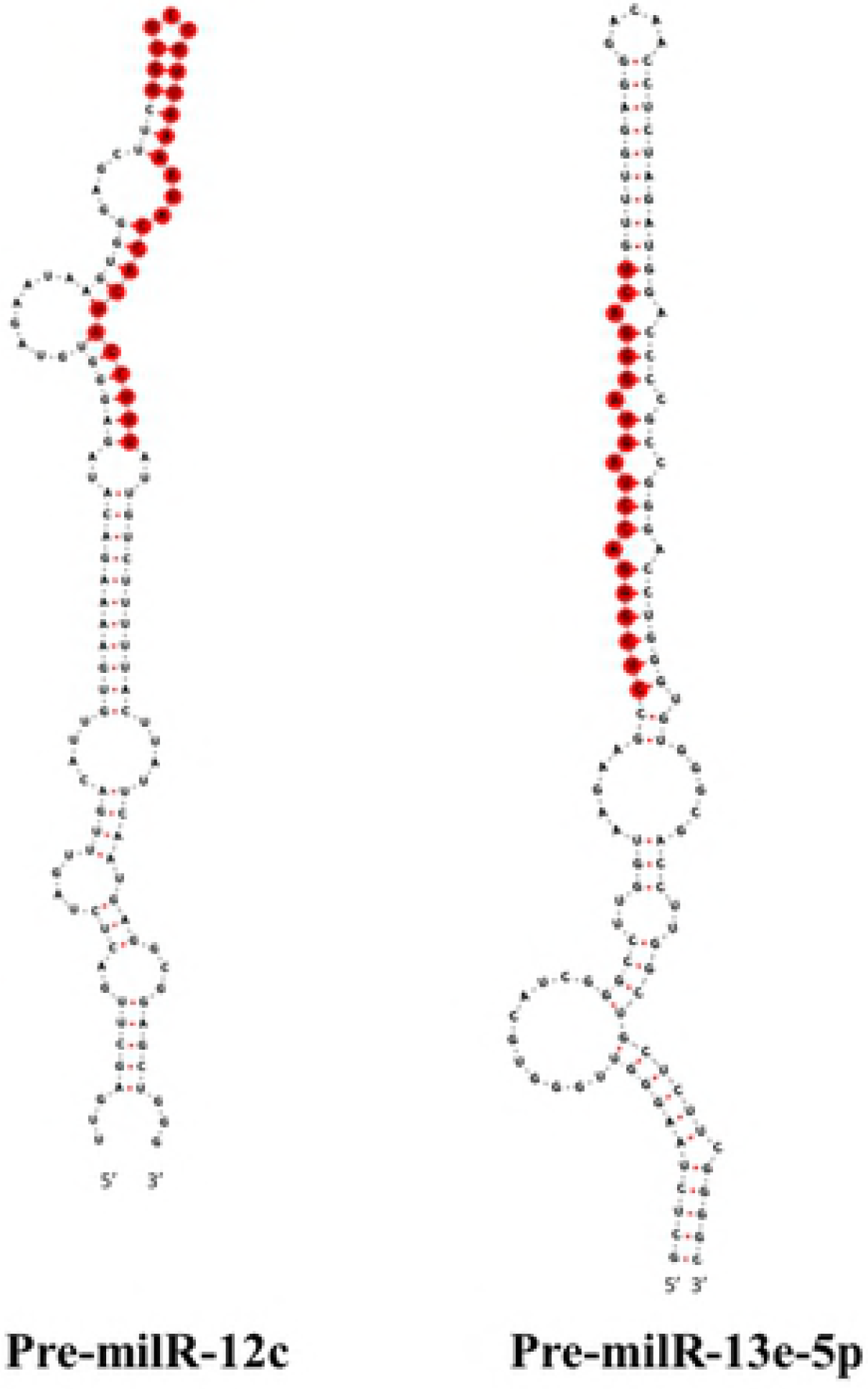
GO annotation of the predicted milRNA targets. The distribution of GO annotation of (a) cci-milR-12c and (b) cci-milR-13e-5p into three major categories. (c) Distribution of detailed GO annotation within each category.

**Fig 11.**
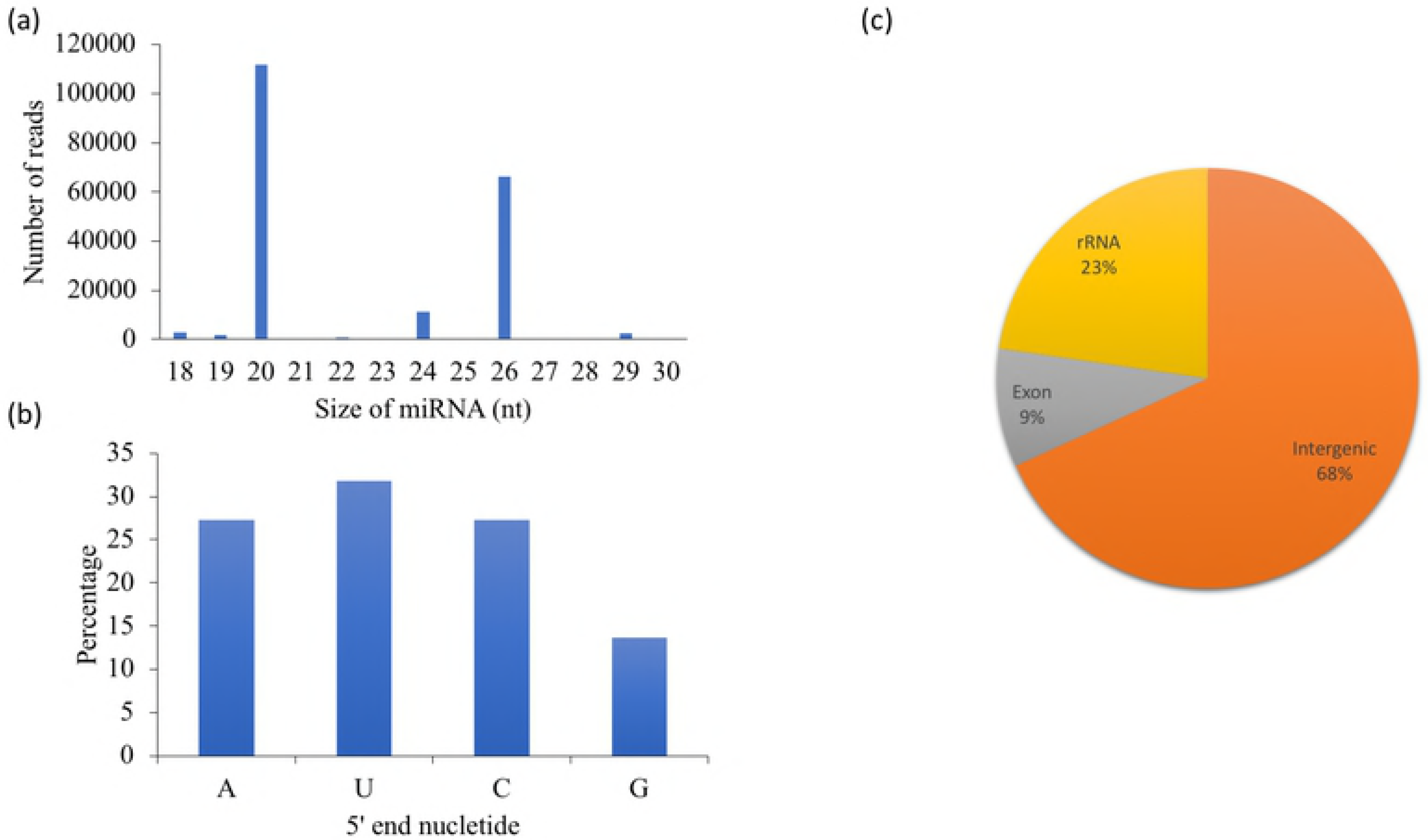
KOG classification of the predicted milRNA targets. Among the annotated KOG terms, “signal transduction mechanisms” (12%), “posttranslational modification, protein turnover, chaperones” (12%), “translation, ribosomal structure and biogenesis” (11%), and “RNA processing and modification” (7%) were the major subcategories of cci-milR-13e-5p putative targets (Fig 11). Overall, results from the GO and KOG term annotations of the two validated milRNAs were similar. None of the targets of cci-milR-12c was assigned to the KEGG pathways and putative targets of cci-milR-13e-5p were annotated to 130 different pathways, most of them were classified into “metabolic pathways”, “biosynthesis of secondary metabolites”, “carbon metabolism”, “oxidative phosphorylation”, “RNA transport”, and “Tight junction” (Fig 12).

**Fig 12.**
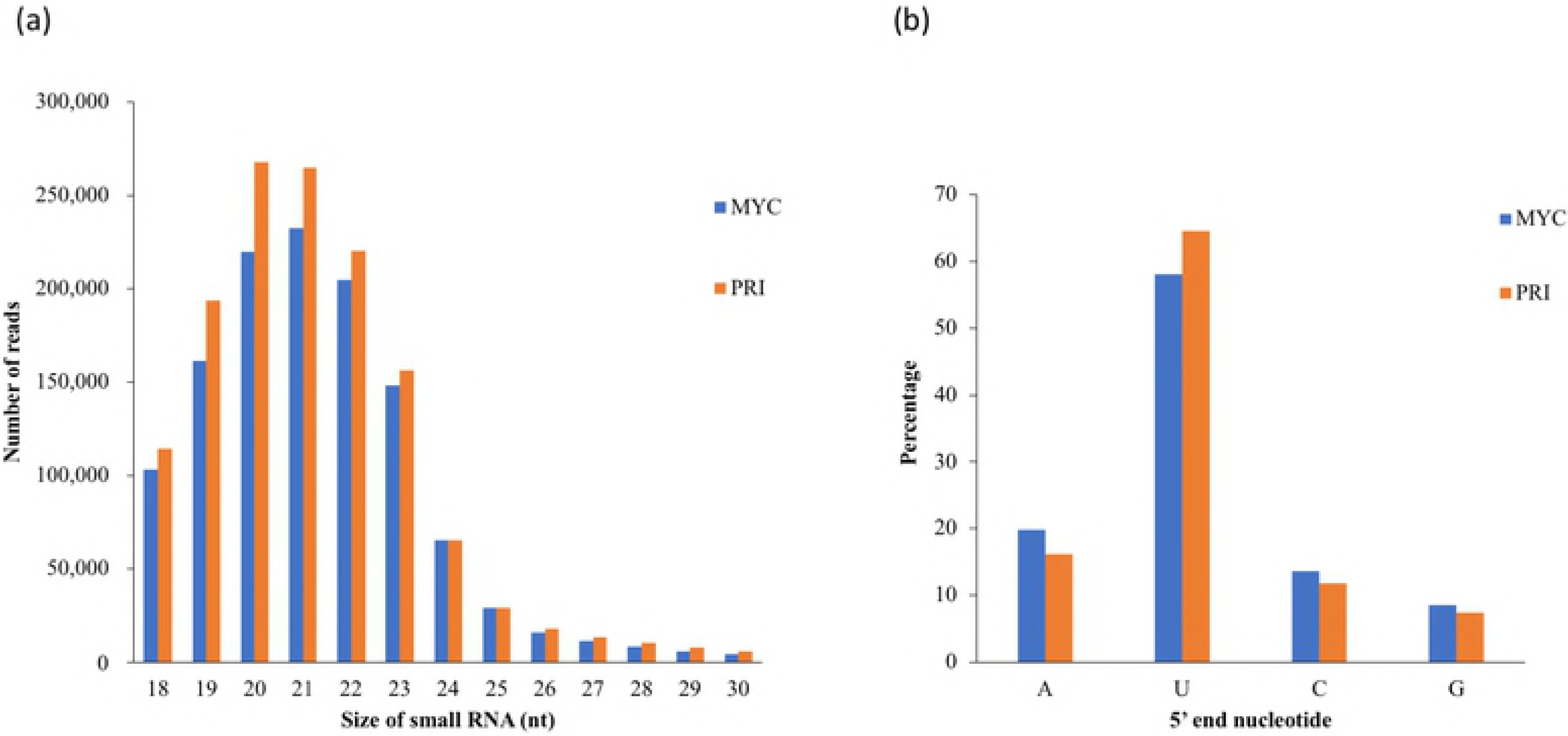
KEGG analysis of predicted cci-milR-13e-5p targets. KEGG biological pathways of 204 predicted cci-milR-13e-5p targets were assigned using the KEGG KAAS with the BBH method and all fungal species were selected as the representative gene set.

Interestingly, both validated milRNAs targeted several fruiting body related genes (S3 Table). Cci-milR-12c targeted a fungal pheromone, hydrophobin and cytochrome P450. Cci-milR-13e-5p targeted protein kinases, transcription factors, heat shock protein, actin, cytochrome P450, and genes related to nucleic acid processing and cell cycle control. Altogether, these results suggest that milRNAs may play an important role for controlling the differential gene expressions and facilitating the cellular developments during the early developmental transition in *C. cinerea*.

## Discussion

In this study, we constructed sRNA libraries and identified milRNAs of *C. cinerea* at two different developmental stages. Characteristics of *C. cinerea* milRNA populations similar to those in animals and plants and the presence of core proteins of miRNA biogenesis in *C. cinerea* suggest that milRNAs in mushrooms may be produced in similar pathways to those in animals and plants. Moreover, the functional analysis of milRNA targets demonstrates the potential regulatory roles of milRNAs in fruiting body development (S3 Fig).

The expression patterns of milRNAs give a hint on their biological functions across different biological processes. Here, we identified putative targets that exhibited a negative correlation in expression profiles with milRNAs during the transition from MYC to PRI, some of which were related to fruiting body formation. cci-milR-12c potentially controls mycelial development by targeting fungal pheromone and hydrophobin. Fungal pheromone is responsible for initiating septal dissolution and clamp-cell fusion during the transition from monokaryotic to dikaryotic state, whereas hydrophobin regulates morphogenesis in fungi, particularly in fruiting body development of basidiomycetes [22, 67, 68]. Besides, it has been reported that different sets of hydrophobins are employed by mushroom forming basidiomycetes in different developmental stages [68, 69]. On the contrary, cci-milR-13e-5p targeted protein kinases, cytochrome P450, laccase, actin, and genes related to nucleic acid processing. Protein kinase has been predicted to respond to nutrient depletion in fruiting body initiation, especially FunK1, which can only be found in multicellular fungi. In addition, the up-regulated heat shock proteins in PRI are response to the lower temperature for fruiting body development (25°C) than that of mycelial cultivation (37 °C) [18, 19, 70–73]. Therefore, cci-milR-13e-5p is more likely to regulate cellular metabolism during the early development transition, for instance, by supporting the dynamic structural changes and regulating oxidative phosphorylation, carbon metabolism and nucleotide metabolism, in response to an increased demand of DNA synthesis, energy production, protein synthesis and turnover from the mycelial to primordium stage [24]. Results suggest that milRNAs may play a role in controlling the drastic transcriptomic and morphological changes during fruiting body initiation.

Phylogenetic analysis results of DCLs, AGO-like and QDE-2 proteins and the fact that no miRNA homologs of animals and plants were identified in the *C. cinerea* genome, providing evidence to the claim that miRNAs may evolve independently among animals, plants and fungi [31, 74, 75, 76]. Our results also indicate an early duplication and diversification of Dicer proteins followed by a lineage-specific loss of PAZ domain in fungi. DCL-1 (CC1G_00230) is evolutionary closely related to other mushroom forming fungi and is the only DCL in *C. cinerea* that contains the PAZ domain, which has only been found in mushroom forming basidiomycetes. The PAZ domain recognizes the 3’ 2-nt overhang of pre-miRNA during miRNA biogenesis and the specific distance between the anchoring site of PAZ and RNase III domain is used to determine the milRNA product size [77]. However, this functional domain is absent in other fungal species, suggesting that different molecular mechanisms are adopted by DCLs without the PAZ domain to produce milRNAs with heterogeneity in length. Indeed, size heterogeneity of fungal milRNAs has been reported in *N. crassa* and *F. oxysporum* [11, 12]. Besides, homologs of cci-milR-12c were found in another mushroom, *L. bicolor*. These results suggest that milRNAs are produced in an alternative way among mushroom forming fungi and serve similar functions to regulate the development of multicellular structures in fungi.

Given that miRNAs are generally produced from a hairpin precursor by Dicer, the accumulation of pre-miRNAs can be detected in organisms with impaired Dicer function [1]. Change of miRNA expression patterns is an indicator of the participation of Dicer in its biogenesis. Although homologous recombination has been found in *C. cinerea*, gene knockouts are difficult to achieve due to the high efficiency of non-homologous DNA end joining [78]. Therefore, an alternative gene silencing method, dsRNA-mediated gene knockdown, which was successfully used in the study of *C. cinerea* strains #326 *(A43mut B43mutpab1-1),* was used in this study [79]. However, cci-milR-12c and cci-milR-13e-5p were still produced - corresponding RNA bands of these milRNAs were detected in northern blot, with a ~70% knockdown efficiency of DCLs. It is possible that milRNAs are efficiently produced, even when the expression levels of the DCLs are extremely low. Future works are needed to investigate the roles of PAZ-containing DCLs in milRNA biogenesis of mushrooms and to determine if the milRNA homologs play a regulatory role in other mushroom forming fungi.

## Conclusions

Our findings have demonstrated differential post-transcriptional regulatory roles of milRNAs in different developmental stages of the mushroom forming fungus *C. cinerea* and identified the milRNA potential targets involved in fruiting body formation, providing new insights into the regulatory mechanisms of fruiting body development and the potential functions of milRNAs in fungi. Moreover, we have found putative core miRNA biogenesis proteins, Dicer and AGO, in the *C. cinerea* genome. Phylogenetic analysis showed that these proteins were more closely related to those in other fungal species than in animals and plants. However, the roles of DCLs, AGO-like and QDE-2 proteins in the biogenesis of *C. cinerea* milRNAs cannot be identified here. Altogether, these results serve as the foundation for further evolutionary developmental studies of fungi and contribute to the phylogenetic occurrence of miRNA-mediated regulatory system among different kingdoms.

## Acknowledgements

We thank Dr. Hajime Muraguchi for sharing *C. cinerea* strains used in this study, Dr. Yi Liu for his insightful discussions on milRNAs and Mr. Tsz Kai Li for technical assistance.

## Supporting information

**S1 Table. Primers used in RT-qPCR for expression determination of core biogenesis proteins.**

F: forward primer, R: reverse primer.

**S2 Table. Stealth siRNA duplexes used in DCLs knockdown assays.**

S: sense strand, AS: antisense strand of siRNA duplexes.

**S3 Table. Target prediction of two validated milRNAs in *C. cinerea.***

Predicted targets of (a) cci-milR-12c and (b) cci-milR-13e-5p. Norm_MYC and Norm_PRI represent normalized expression levels at the mycelium (MYC) and primordium (PRI) stages based on previously published microarray data of *C. cinerea* [25]. Targets were predicted by using miRanda, PITA and microTar and selected by several rounds of functional annotation. Description and domain information are downloaded from http://www.broadinstitute.org.

**S1 Fig. Schematic 2D domain architecture of Dicer and AGO proteins in *N. crassa.***

The grey bars represent the full protein sequences and the colored boxes represent identified functional domains.

**S2 Fig. Effect of DCL knockdown on miRNA expression.**

(a) RT-qPCR expression levels of DCLs obtained in DCL knockdown strains after normalization against the untreated primordium (control). Results were obtained from three independent experimental replicates. The treatment samples were significantly different from the control samples. *p < 0.05, ** p < 0.01. Northern blot of sRNA samples shows the presence of (b) cci-milR-12c, (c) cci-milR-13e-5p and their precursors in all the knockdown strains. The top panels show the northern blots probed with milRNA-specific DIG probes. The 15% denaturing gels stained with ethidium bromide (EtBr) in the bottom panels indicate equal loading of RNA samples.

**S3 Fig. Schematic summary of the milRNA study in *C. cinerea.***

## References

1. Ambros V, Bartel B, Bartel DP, Burge CB, Carrington JC, Chen X, et al. A Uniform System for microRNA Annotation. RNA. 2003 Mar 9;9;277–277. pmid: 12592000.

2. Kutter C, Svoboda P. miRNA, siRNA, piRNA: Knowns of the unknown. RNA biol. 2008 Oct-Dec; 5(4):181. pmid: 19182524.

3. Tili E, Michaille JJ, Calin GA. Expression and function of microRNAs in immune cells during normal or disease state. Int J Med Sci. 2008 Apr 3;5(2):73–79. pmid: 18392144.

4. Zhang B, Pan X, Cobb GP, Anderson TA. Plant microRNA: A small regulatory molecule with big impact. Dev Biol. 2006 Jan 1;289;3–16. pmid: 16325172.

5. Banerjee D, Slack F. Control of developmental timing by small temporal RNAs: a paradigm for RNA-mediated regulation of gene expression. Bioessays. 2002 Feb 24; 24(2): 119–29. pmid: 11835276.

6. Brennecke J, Hipfner DR, Stark A, Russell RB, Cohen SM. bantam encodes a developmentally regulated microRNA that controls cell proliferation and regulates the proapoptotic gene hid in Drosophila. Cell. 2003 Apr 4;113(1):25–36 pmid: 12679032

7. Chen CZ, Schaffert S, Fragoso R, Loh C. Regulation of immune responses and tolerance: the microRNA perspective. Immunol Rev. 2013 May; 253(1): 112–28. pmid: 23550642.

8. Lee RC, Feinbaum RL, Ambros V. The C. elegans heterochronic gene lin-4 encodes small RNAs with antisense complementarity to lin-14. Cell. 1993 Dec 3;75(5):843–54. pmid:8252621

9. Wienholds E, Plasterk RH. MicroRNA function in animal development. FEBS letters. 2005 Oct 31; 579(26):5911–22. pmid: 16111679.

10. Bartel DP. MicroRNAs: Genomics, Biogenesis, Mechanism, and Function. Cell. 2004 Jan 23;116(2):281–97. pmid: 14744438.

11. Lee HC, Li L, Gu W, Xue Z, Crosthwaite SK, Pertsemlidis A, et al. Diverse pathways generate microRNA-like RNAs and Dicer-independent small interfering RNAs in fungi. Mol cell., 2010 Jun 25;38(6):803–14. pmid: 20417140.

12. Chen R, Jiang N, Jiang Q, Sun X, Wang Y, Zhang H, et al. Exploring microRNA-like small RNAs in the filamentous fungus Fusarium oxysporum. PLoS ONE. 2014 Aug 20;9(8): e104956.

13. Kang K, Zhong J, Jiang L, Liu G, Gou CY, Wu Q, et al. Identification of microRNA-Like RNAs in the filamentous fungus Trichoderma reesei by solexa sequencing. PLoS ONE. 2013 Oct 2; 8(10): e76288.

14. Lau SKP, Chow WN, Wong AYP, Yeung JMY, Bao J, Zhang N, et al. Identification of microRNA-like RNAs in mycelial and yeast phases of the thermal dimorphic fungus Penicillium marneffei. PLoS Negl Trop Dis. 2013 Auh 22; 7(8): e2398.

15. Lin YL, Man LT, Lee YR, Lin SS, Wang SY, Chang TT, et al. MicroRNA-like small RNAs prediction in the development of Antrodia cinnamomea. PLoS ONE. 2015 Apr 10;10(4): e0123245.

16. Zhou J, Fu Y, Xie J, Li B, Jiang D, Li G, et al. Identification of microRNA-like RNAs in a plant pathogenic fungus Sclerotinia sclerotiorum by high-throughput sequencing. Mol Genet Genomics. 2012 Apr;287(4):275–82. pmid: 22314800.

17. Stajich JE, Berbee ML, Blackwell M, Hibbett DS, James TY, Spatafora JW, et al. Primer-The fungi. Curr Biol. 2009 Sep 29;19(18):R840–R845. pmid: 19788875.

18. Stajich JE, Wilke SK, Ahrén D, Au CH, Birren BW, Borodovsky M, et al. Insights into evolution of multicellular fungi from the assembled chromosomes of the mushroom Coprinopsis cinerea (Coprinus cinereus). Proc Natl Acad Sci U S A. 2010 Jun 29;107(26):11889–94. pmid: 20547848.

19. Moore D. Graviresponses in fungi. Adv Space Res. 1996;17(6-7):73–82.

20. Terashima K, Yuki K, Muraguchi H, Akiyama M, Kamada T. The dst1 gene involved in mushroom photomorphogenesis of Coprinus cinereus encodes a putative photoreceptor for blue light. Genetics. 2005 Sep;171(1):101–108. pmid: 15956671.

21. Kuratani M, Tanaka K, Terashima K, Muraguchi H, Nakazawa T, Nakahori K, et al. The dst2 gene essential for photomorphogenesis of Coprinopsis cinerea encodes a protein with a putative FAD-binding-4 domain. Fungal Genet Biol. 2010 Feb 4;47(2):152–8. pmid: 19850145.

22. Kües U. Life history and development processes in the basidiomycete Coprinus cinereus. Microbiol Mol Biol Res. 2000 Jun;64(2):316–53. pmid:.10839819.

23. Kamada T, Sano H, Nakazawa T, Nakahori K. Regulation of fruiting body photomorphogenesis in Coprinopsis cinerea. Fungal Genet Biol. 2010 Nov;47(11):917–21. pmid: 20471485.

24. Cheng CK, Au CH, Wilke SK, Stajich JE, Zolan ME, Pukkila PJ, et al. 5′-Serial Analysis of Gene Expression studies reveal a transcriptomic switch during fruiting body development in Coprinopsis cinerea. BMC genomics. 2013 Mar 20;14(1):195.

25. Cheng X, Hui JH, Lee YY, Wan Law PT, Kwan HS. A “developmental hourglass” in fungi. Mol Biol Evol. 2015 Jun;32(6):1556–66. pmid: 25725429.

26. Berezikov E, Guryev V, van de Belt J, Wienholds E, Plasterk RH, Cuppen E. Phylogenetic shadowing and computational identification of human microRNA genes. Cell. 2005 Jan 14;120(1):21–4. pmid: 15652478.

27. Lau NC, Lim LP, Weinstein EG, D.P. Bartel. An abundant class of tiny RNAs with probable regulatory roles in Caenorhabditis elegans. Science. 2001 Oct 26;294(5543):858–62. pmid: 11679671.

28. Lagos-Quintana M, Rauhut R, Lendeckel W, Tuschl T. Identification of novel genes coding for small expressed RNAs. Science. 2001 Oct 26;294(5543):853–8. pmid: 11679670.

29. Lee RC, Ambros V. An extensive class of small RNAs in Caenorhabditis elegans. Science. 2001 Oct 26;294(5543):962–4. pmid: 11679672.

30. Xie X, Lu J, Kulbokas EJ, Golub TR, Mootha V, Lindblad-Toh K, et al. Systematic discovery of regulatory motifs in human promoters and 3′ UTRs by comparison of several mammals. Nature. 2005 Mar 17;434(7031):338–45. pmid: 15735639.

31. Yang Q, Li L, Xue Z, Ye Q, Zhang L, Li S, et al. Transcription of the Major Neurospora crassa microRNA-Like Small RNAs Relies on RNA Polymerase III. PLoS Genet. 2013 Jan 17;9(1):e1003227.

32. Houbaviy HB, Murray MF, Sharp PA. Embryonic stem cell-specific microRNAs. Dev. Cell. 2003 Aug;5(2):351–8. pmid: 12919684.

33. Lagos-Quintana M, Rauhut R, Yalcin A, Meyer J, Lendeckel W, Tuschl T. Identification of tissue-specific microRNAs from mouse. Curr. Biol. 2002 Apr 30;12(9):735–9. pmid: 12007417.

34. Shamimuzzaman, M, Vodkin L. Identification of soybean seed developmental stage-specific and tissue-specific miRNA targets by degradome sequencing. BMC genomics. 2012 July 16; 13(1):310.

35. Watanabe T, Takeda A, Mise K, Okuno T, Suzuki T, Minami N, et al. Stage-specific expression of microRNAs during Xenopus development. FEBS Lett. 2005 Jan 17;579(2):318–24. pmid: 15642338.

36. Valentine G, Wallace YJ, Turner FR, Zolan ME. Pathway analysis of radiation-sensitive meiotic mutants of Coprinus cinereus. Mol Gen Genet. 1995 Apr 20;247(2):169–79. pmid: 7753026.

37. Muraguchi H, Umezawa K, Niikura M, Yoshida M, Kozaki T, Ishii K, et al. Strand-specific RNA-seq analyses of fruiting body development in Coprinopsis cinerea. PLoS ONE. 2015 Oct 28;10(10):e0141586.

38. Burns C1, Stajich JE, Rechtsteiner A, Casselton L, Hanlon SE, Wilke SK, et al. Analysis of the Basidiomycete Coprinopsis cinerea reveals conservation of the core meiotic expression program over half a billion years of evolution. PLoS Genet. 2010 Sep 23;6(9):e1001135. doi: 10.1371/journal.pgen.1001135.

39. Srivilai P, Loutchanwoot P. Coprinopsis cinerea as a model fungus to evaluate genes underlying sexual development in basidiomycetes. Pak J Biol Sci. 2009 Jun 1;12(11):821–35. pmid: 19803116

40. Gardner PP, Daub J, Tate JG, Nawrocki EP, Kolbe DL, Lindgreen S, et al. (2009). Rfam: updates to the RNA families database. Nucl Acids Res. 2009 Jan;37(Database issue):D136–40. doi: 10.1093/nar/gkn766.

41. Langmead B, Trapnell C, Pop M, Salzberg SL. Ultrafast and memory-efficient alignment of short DNA sequences to the human genome. Genome biol. 2009;10(3):R25. pmid: 19261174.

42. Kozomara A, Griffiths-Jones S. miRBase: annotating high confidence microRNAs using deep sequencing data. Nucl Acids Res. 2014 Jan;42(Database issue):D68–73. pmid: 24275495

43. Bonnet E, Wuyts J, Rouzé P, Van de Peer Y. Evidence that microRNA precursors, unlike other non-coding RNAs, have lower folding free energies than random sequences. Bioinformatics. 2004 Nov 22;20(17):2911–7. pmid: 15217813.

44. Hofacker IL, Fontana W, Stadler PF, Bonhoeffer LS, Tacker M, Schuster P. Fast Folding and Comparison of RNA Secondary Structures. Monatshefte für Chemie/Chemical Monthly. 1994 Feb;125(2):167–88.

45. Lorenz R, Bernhart SH, Höner Zu Siederdissen C, Tafer H, Flamm C, Stadler PF, et al. ViennaRNA package 2.0. Algorithms Mol Biol. 2011 Nov 24;6:26. pmid: 22115189.

46. Kim SW, Li Z, Moore PS, Monaghan AP, Chang Y, Nichols M, et al. A sensitive non-radioactive northern blot method to detect small RNAs. Nucl Acids Res. 2010 Apr;38(7):e98. pmid: 20081203.

47. Robert D Finn, Alex Bateman, Jody Clements, Penelope Coggill, Ruth Y. Eberhardt, Sean R. Eddy, et al. Pfam: the protein families database. Nucl Acids Res. 2014 Jan 1; 42(Database issue): D222–D230. pmid: 24288371.

48. Schultz J1, Copley RR, Doerks T, Ponting CP, Bork P. SMART: a web-based tool for the study of genetically mobile domains. Nucl Acids Res. 2000 Jan 1;28(1):231–4. pmid: 10592234.

49. Kumar S, Stecher G, and Tamura K. MEGA7: Molecular Evolutionary Genetics Analysis version 7.0 for bigger datasets. Mol Biol Evol. 2016 Jul;33(7):1870–4. pmid: 27004904.

50. Kramer MF. Stem-Loop RT-qPCR for miRNAs. Curr Protoc Mol Biol. 2011 Jul;15(10). pmid: 21732315.

51. Holmes K, Williams CM, Chapman EA, Cross MJ. Detection of siRNA induced mRNA silencing by RT-qPCR: considerations for experimental design. BMC Res Notes. 2010 Mar 3;3:53. pmid: 20199660.

52. Enright AJ1, John B, Gaul U, Tuschl T, Sander C, Marks DS. MicroRNA targets in Drosophila. Genome Biol. 2003;5(1):R1. pmid: 14709173.

53. Kertesz M, Iovino N, Unnerstall U, Gaul U, Segal E. The role of site accessibility in microRNA target recognition. Nat. Genet. 2007 Sep 23;39:1278–84.

54. Thadani R, Tammi MT. MicroTar: predicting microRNA targets from RNA duplexes. BMC Bioinformatics. 2006;7(Suppl 5):S20. pmid: 17254305.

55. Zhang Y, Verbeek FJ. Comparison and integration of target prediction algorithms for microRNA studies. J. Integr. Bioinform. 2010 Mar 25;7(3). pmid: 20375447.

56. Tatusov RL, Fedorova ND, Jackson JD, Jacobs AR, Kiryutin B, Koonin EV, et al. The COG database: an updated version includes eukaryotes. BMC Bioinformatics. 2003 Sep 11;4:41. pmid: 12969510.

57. Götz S, García-Gómez JM, Terol J, Williams TD, Nagaraj SH, Nueda MJ, et al. High-throughput functional annotation and data mining with the Blast2GO suite. Nucl Acids Res. 2008 Jun;36(10):3420–35. pmid: 18445632.

58. Moriya Y, Itoh M, Okuda S, Yoshizawa AC, Kanehisa M. KAAS: an automatic genome annotation and pathway reconstruction server. Nucl Acids Res. 2007 Jul;35(Web Server issue):W182–185. pmid: 17526522.

59. Stajich JE, Wilke SK, Ahrén D, Au CH, Birren BW, Borodovsky M, et al. Coprinopsis cinerea Genome Project; 2010 [cited 2010 Jun 29]. Database: Broad Institute [Internet]. Available from: http://www.broadinstitute.org/annotation/genome/coprinus_cinereus/MultiHome.html

60. Zhang Y, Zhang R, Su B. Diversity and evolution of MicroRNA gene clusters. Science in China. Science in China Series C: Life Sciences. 2009 Mar;52(3):261–6.

61. Martin F, Aerts A, Ahren D, Brun A, Danchin EG, Duchaussoy F, et al. The genome of Laccaria bicolor provides insights into mycorrhizal symbiosis. Nature. 2008 Mar 6;452(7183):88–92. doi: 10.1038/nature06556.

62. Zhu QW, Luo YP. Identification of miRNAs and their targets in tea (Camellia sinensis). JJ Zhejiang Univ Sci B. 2013 Oct;14(10):916–23. pmid: 24101208

63. Yoshikawa M, Fujii YR. Human ribosomal RNA-derived resident microRNAs as the transmitter of information upon the cytoplasmic cancer stress. Biomed Res Int. 2016;2016:7562085. pmid: 27517048.

64. Carthew RW, Sontheimer EJ. Origins and mechanisms of miRNAs and siRNAs. Cell. 2009 Feb 20;136(4):642–55. pmid: 19239886.

65. Lau PW, Guiley KZ, De N, Potter CS, Carragher B, MacRae IJ. The molecular architecture of human Dicer. Nat. Struct. Mol. Biol. 2012 Mar 18;19(4):436–40. pmid: 22426548.

66. Höck J, Meister G. The Argonaute protein family. Genome Biol. 2008;9(2):210. pmid: 18304383.

67. Wösten HA. Hydrophobins: multipurpose proteins. Annu Rev Microbiol. 2001;55;625–46. pmid: 11544369.

68. Chum WW, Ng KT, Shih RS, Au CH, Kwan HS. Gene expression studies of the dikaryotic mycelium and primordium of Lentinula edodes by serial analysis of gene expression. Mycol Res. 2008 Aug;112(Pt 8):950–64. pmid: 18555678.

69. Ma A, Shan L, Wang N, Zheng L, Chen L, Xie B. Characterization of a Pleurotus ostreatus fruiting body-specific hydrophobin gene. Po.hyd. J Basic Microbiol. 2007 Aug;47(4):317–24. pmid: 17647210.

70. D’Souza CA, Heitman J. Conserved cAMP signaling cascades regulate fungal development and virulence. FEMS microbiol rev. 2001 May;25(3):349–64. pmid: 11348689.

71. Palmer GE, Horton JS. Mushrooms by magic: making connections between signal transduction and fruiting body development in the basidiomycete fungus Schizophyllum commune. FEMS Microbiol Lett. 2006 Sep; 262(1):1–8. pmid: 16907732.

72. Liu Y, Srivilai P, Loos S, Aebi M, Kües U. An essential gene for fruiting body initiation in the basidiomycete Coprinopsis cinerea is homologous to bacterial cyclopropane fatty acid synthase genes. Genetics. 2006 Feb;172(2):873–84. pmid: 16322509.

73. Robert JC, & Durand, R. Light and temperature requirements during fruit-body development of a basidiomycete mushroom., Coprinus congregates. Physiol Plant. 1979 Jun;46(2):174–8.

74. Moran Y, Agron M, Praher D, Technau U. The evolutionary origin of plant and animal microRNAs. Nat Ecol Evol. 2017 Feb 21;1(3):27. pmid: 28529980.

75. Murphy D, Dancis B, Brown JR. The evolution of core proteins involved in microRNA biogenesis. BMC Evol Biol. 2008 Mar 25;8:92. pmid: PMC2287173.

76. Ruiz-Trillo I, Burger G, Holland PW, King N, Lang BF, Roger AJ, Gray MW. The origins of multicellularity: a multi-taxon genome initiative. Trends Genet. 2007 Mar;23(3):113–8. pmid: 17275133.

77. Kandasamy SK, Fukunaga R. Phosphate-binding pocket in Dicer-2 PAZ domain for high-fidelity siRNA production. Proc Natl Acad Sci U S A. 2016 Dec 6;113(49): 14031–14036. pmid: 27872309.

78. Ninomiya Y, Suzuki K, Ishii C, Inoue H. Highly efficient gene replacements in Neurospora strains deficient for nonhomologous end-joining. Proc Natl Acad Sci USA. 2004 Aug 17;101(33):12248–53. pmid: 15299145.

79. Namekawa SH, Iwabata K, Sugawara H, Hamada FN, Koshiyama A, Chiku H, et al. Knockdown of LIM15/DMC1 in the mushroom Coprinus cinerus by double-sranded RAN-mediated gene silencing. Microbiology. 2005 Nov;151(Pt 11):3669–78. pmid: 16272388.

